# Immunoglobulin sub-class levels define inter-donor plasma variability: a longitudinal dual-lab study

**DOI:** 10.1101/2025.11.04.686385

**Authors:** Stephan Michalik, Sofia Kalaidopoulou Nteak, Nicolas Drouin, Manuela Gesell Salazar, Elke Hammer, Vishnu Mukund Dhople, Silva Holtfreter, Stefan Weiss, Henk van den Toorn, Barbara Bröker, Grażyna Domańska, Uwe Völker, Albert J. R. Heck

## Abstract

Advancements in mass spectrometry have transformed plasma proteomics, allowing for high-throughput analysis of large cohorts. This study utilized the TIMES cohort, consisting of 51 healthy participants monitored monthly over 12 months, to evaluate intra- and inter-individual variability in the plasma proteome. Approximately 600 samples were analyzed in two independent laboratories, revealing strong correlations despite methodological differences. The study focused on the stability of the plasma proteome within donors over a year, with larger differences observed between donors. A support vector machine model achieved a 98% median classification accuracy, confirming stable, donor-specific proteomic profiles. Immunoglobulins, often overlooked in plasma proteomics, were found to contribute significantly to donor-specific signatures, showing pronounced inter-individual variability but remarkable intra-donor stability. In contrast, C-reactive protein and other inflammation markers exhibited significant temporal fluctuations and donor-specific baseline levels. This inter-laboratory study highlights the importance of longitudinal sampling for biomarker discovery, the robustness of MS-based proteomic workflows, and provides new insights into immunoglobulin dynamics and variability in human plasma.

**Synopsis:** 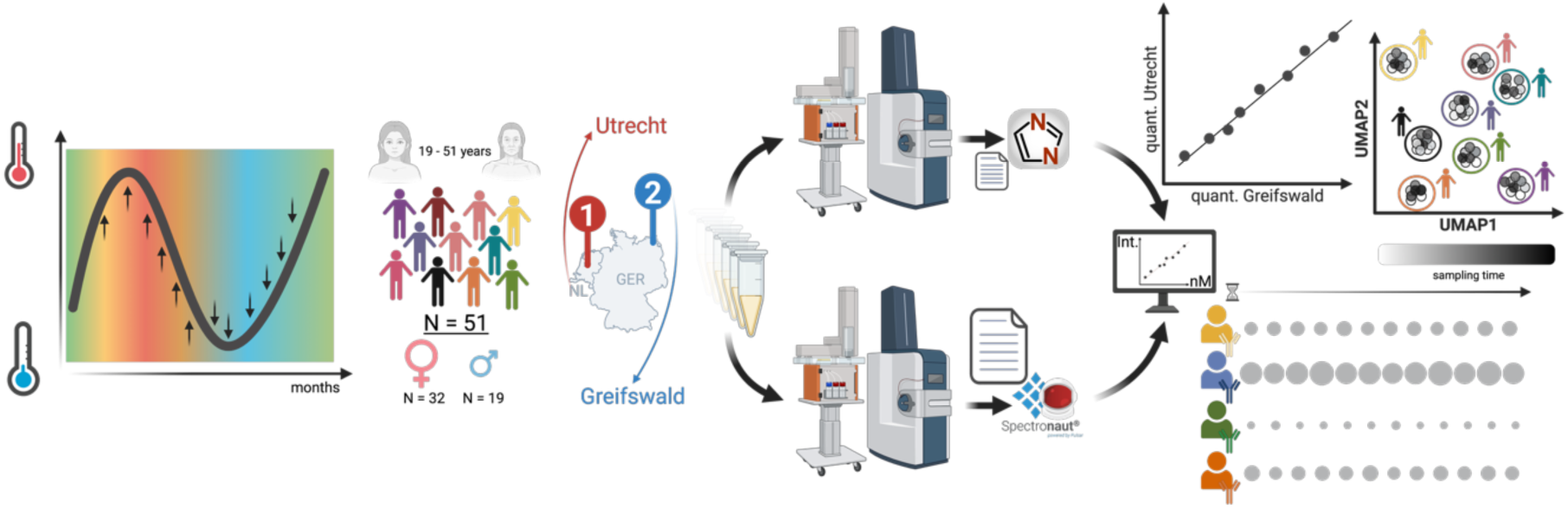

- Longitudinal analysis of individual plasma proteomes over a one-year period revealed low intra-individual variability albeit high inter-individual differences.
- Cross-site laboratory comparisons confirmed strong concordance in analytical performance and measurement consistency.
- Conversion of relative protein intensities to absolute concentrations revealed immunoglobulin levels on par with established clinical reference ranges.
- Donors were distinguishable based on their unique plasma proteome.
- The immunoglobulin sub-classes are key discriminators defining inter- and intra-donor variability

## Introduction

Due to the relatively non-invasive nature of sampling, blood plasma plays a central role in clinical diagnostics and therapeutic monitoring (Geyer *et al*, 2017; Álvez *et al*, 2025). However, the plasma proteome represents one of the analytically most demanding proteomes, characterized by its vast protein diversity and extended dynamic range (Anderson, 2002). The inherent complexity of the plasma proteome reflects its multifaceted biological origins. Circulating proteins originate from multiple sources: hepatocytes secrete the majority of plasma proteins, lymphocytes produce immunoglobulins and cytokines, or proteins entering the circulation through tissue-damage-induced leakage (Zhang *et al*, 2007). This complexity is further amplified by the polymorphic nature of several plasma proteins, most notably the immunoglobulins, as well as the extensive array of post-translational protein modifications (PTMs), especially glycosylation (Nimmerjahn *et al*, 2023). Furthermore, concentrations of plasma proteins span a range of over ten orders of magnitude, e.g., from highly abundant proteins such as albumin, accounting for 55% of the total protein content (Anderson, 2002), and immunoglobulins (about 20% (Patel *et al*, 2023)), down to trace-level cytokines (Geyer *et al*, 2017; Ameling *et al*, 2025).

Traditionally, blood protein analysis relies on targeted techniques such as ELISA, which enable robust quantification of predefined protein targets within complex mixtures. In contrast, mass spectrometry (MS) offers a powerful alternative for both diagnostic applications and biomarker discovery, as protein identification, quantification and characterization are performed unbiased (Aebersold & Mann, 2016). Recent advancements in MS have improved the analytical capabilities of plasma proteomics. Enhanced sensitivity, expanded dynamic range of mass spectrometry devices (Vitko *et al*, 2024; Hendricks *et al*, 2024), and refined sample preparation protocols now enable the differentiation of physiological or pathological states, which is essential for disease screening, diagnosis, therapeutic stratification, and monitoring treatment efficacy. Despite substantial investment in biomarker research, the clinical translation of novel protein biomarkers remains limited. While FDA approvals have incorporated an increasing number of distinct cancer biomarkers, growing from 3 in 2011 to 16 in 2022 (Dou *et al*, 2024), the pace of new biomarker adoption remains modest relative to the scale of research investment. A major barrier to clinical translation is the lack of standardization across the biomarker discovery pipeline: challenges in sample collection, pre-analytical handling, measurement platforms, and data processing hinder cross-study integration and limit the generation of robust, comparable results across laboratories, cohorts, and populations (Cai *et al*, 2025). These translational bottlenecks highlight the necessity for a robust and integrated screening pipeline and workflows for biomarker discovery research and clinical proteomics (Völlmy *et al*, 2021; Messner *et al*, 2020). Recent comparative analyses of 16 distinct proteomic sample preparation methodologies (Varnavides *et al*, 2022), evaluated in HeLa cell lysates, revealed that most preparation protocols yielded comparable protein extraction efficiencies, demonstrated high inter-method reproducibility, and exhibited minimal levels of preparation-induced artifacts. These findings emphasize that, despite the inherent methodological variability in proteomics sample preparation and analytical workflows, harmonization is achievable and can yield consistent results across laboratories and studies.

Here, we set out to empirically validate this concept of harmonization making use of the TIMES (Tracking Individuals Monthly for Evaluating Stability) cohort. This cohort comprises of 51 (supposedly) healthy participants from whom blood was retrieved monthly over a period of 12 months, resulting in about ∼600 samples. This allowed us to assess intra- and inter-individual variability in the plasma proteome in a healthy cohort. We independently processed and analyzed the healthy donor cohort at two separate laboratories in Germany and the Netherlands employing distinct sample preparation protocols, mass spectrometry platforms and initial raw data analysis workflows.

## Materials & Methods

### Study design and participants

The TIMES study aimed to investigate the circannual fluctuations in the plasma proteome of healthy donors over a period of 12 months. Inclusion criteria included (i) age 18 - 60 years, (ii) written informed consent, and (iii) no acute illness in the last seven days before sampling. Exclusion criteria included (i) body mass index < 18.5 kg/m^2^, and (ii) presence of blood coagulation disorders, anemia or similar conditions. Plasma samples were obtained from a total of 53 participants once per month for one year (June 2017 – May 2018). Two subjects were excluded from data analyses due to early loss to follow-up. Data from the remaining 51 subjects were included in the analyses (Supplemental Table 1).

### Ethics and data protection

The TIMES study was approved by the Ethics Committee of the University Medicine Greifswald (internal registration numbers BB 043/17 and BB 001/21f). All work was conducted in accordance with the tenets of the Declaration of Helsinki (version 2013, Fortaleza). All requirements of data protection and confidentiality according to local regulations, the State Data Protection Act Mecklenburg-Western Pomerania, the European Data Protection Directive 95/46/EC, and the General Data Protection Regulation (GDPR) were fully met.

### Plasma sample collection

Blood samples were collected using EDTA-Vacutainer (BD, K2E RAF 368861) tubes, with BD Vacutainer Safety Lock needles (REF 367282) facilitating the draw. Post-collection, the samples were maintained at room temperature for a maximum of one hour prior to centrifugation. Centrifugation parameters were meticulously controlled, with plasma samples subjected to 2000 x g for 10 minutes at room temperature. Plasma extraction was performed carefully, leaving approximately 0.5 cm of plasma above the PBMC layer to maintain sample integrity. All aliquots were promptly stored at −80°C. The timeframe from blood collection, which commenced between 7:30 and 9:00 a.m. during the University Medicine Greifswald (UMG) blood donation session, to the freezing of aliquots did not exceed 11:00 a.m. Samples were processed concurrently with ongoing collection, with batches of 24 samples transported to the laboratory for immediate mixing and centrifugation, ensuring a maximum pre-centrifugation delay of one hour. Samples were stored at −70°C and shipped on dry-ice to the Utrecht lab site.

### Plasma proteomics – Utrecht site

The plasma proteomics workflow was adapted from previous work (Völlmy *et al*, 2021; Nteak *et al*, 2024). 22 µL of plasma samples were subjected to a 20-fold dilution using 100 mM Tris buffer at pH 8.5. Subsequently, 40 µL of the diluted samples were transferred into 20 µL of Buffer 1, comprising of 4% sodium deoxycholate, 160 mM chloroacetamide, and 100 mM Tris at pH 8.5, followed by the addition of 20 µL Buffer 2, containing 40 mM tris-(2-carboxyethyl)-phosphine in 100 mM Tris at pH 8.5. The mixture was incubated at 95°C for 10 minutes to facilitate denaturation and reduction/alkylation processes. Post-incubation, 100 µL of 100 mM Tris buffer at pH 8.5 was added to dilute the mixture followed by addition of 10 µL each of Trypsin (Sigma-Aldrich) and LysC (Wako), both at a stock concentration of 0.2 µg/µL. The samples were then incubated overnight at 37°C to ensure complete digestion. Enzymatic activity was quenched with the addition of 200 µL of 2% trifluoroacetic acid (TFA). The resulting digest was further diluted 36-fold in 1% TFA, yielding a final concentration of approximately 10 ng/µL (based on an average plasma protein concentration of 70µg/µL). 20 µL of roughly 200 ng of total protein were loaded onto Evosep tips following the manufacturer’s instructions. The samples were analyzed using the 60 SPD method on an Evosep ONE LC (Evosep, Odense, Denmark) with an EV-1109 analytical column (ReproSil Saphir C18, 1.5 μm beads by Dr Maisch; 8 cm length × 150 μm i.d.; Evosep). Gradient elution was achieved using mobile phases A (0.1% HCOOH in Milli-Q water) and B (0.1% HCOOH in 100% CH_3_CN). The LC was coupled to a timsTOF HT mass spectrometer (Bruker Daltonics, Bremen, Germany). Peptide ionization was ensured using a captive source equipped with a ZDV sprayer (20 µm ID, 1865691, Bruker Daltonics). The sprayer was operated at a spray voltage of 1,600 V, with drying gas flowing at 3 l/minute and drying at 180°C. The dia-PASEF-MS mode was configured for MS and ion mobility windows, as described in supplemental table 3.

Peptides were isolated within the m/z range of 300-1200, with 12 dia-PASEF scans and 8.3% duty cycle. The IM range was set to 1.6 and 0.6 V cm^2^ and the accumulation and ramp times were set as 100 ms. Collision energy ramping was performed according to the manufacturer’s default settings (20-59 eV within the mobility range of 0.6-1.6 Vs/cm^2^). Acquisition was conducted with a ramp and accumulation time of 100 ms, achieving a 100% duty cycle.

### Plasma proteomics – Greifswald site

The plasma samples were diluted at a ratio of 1:50 in a solution comprising 50 mM HEPES buffer at pH 8, supplemented with 1% sodium dodecyl sulfate (SDS) to ensure optimal protein solubilization. Subsequently, the diluted samples underwent thermal denaturation at 95°C for a duration of 5 minutes. Upon returning to ambient temperature, the samples were processed through the OT-2 assisted MassSpecPreppy workflow (Reder *et al*, 2023), enabling precise protein concentration determination and enzymatic digestion. Three µg of total protein was reduced with dithiothreitol (DTT) and alkylated using iodoacetamide (IAA). The enzymatic digestion was conducted at an enzyme-to-protein ratio of 1:25. To stop the proteolytic digestion, after at least 16 h 2.44 µl of 5% (v/v) TFA was added to the peptide mixture. The final preparation entailed the loading of 300 ng of the digested material onto EvoTips for subsequent mass spectrometric analysis. The samples were measured with a 60 SPD method using an Evosep performance column (EV-1109, 8 cm × 150 µm × 1.5 µm) on an Evosep ONE LC (Evosep, Odense, Denmark) coupled to a timsTOF Pro 2 (Bruker Daltonics, Bremen, Germany). Peptide ionization was ensured using a captive source equipped with a ZDV sprayer (20µm ID, 1865710, Bruker Daltonics, Bremen, Germany). The sprayer was operated at a spray voltage of 1,640-1740 V, with drying gas flowing at 3 L/min and drying at 180°C. The dia-PASEF-MS mode was configured for MS and ion mobility windows, as described in supplemental table 2.

Peptides were isolated within the m/z range of 330.17-1638.48, comprising 16 m/z windows with two ion mobility steps and eight dia-PASEF scans each. These scans were performed within the ion mobility range of 0.57-1.47 Vs/cm^2^, resulting in a cycle time of 0.95 seconds. Collision energy ramping was performed according to the manufacturer’s default settings (20-52 eV within the mobility range of 0.6-1.6 Vs/cm^2^). Acquisition was conducted with a ramp and accumulation time of 100 ms, achieving a 100% duty cycle.

### Raw data analysis – Greifswald site

Raw data analysis was conducted using Spectronaut® version 20 in directDIA+ mode, employing reviewed Uniprot databases (*Homo sapiens*, comprising 42,529 canonical and isoform entries). Acetylation at the N-terminus of a protein, methionine oxidation were set as variable, and carbamidomethylation on cysteine residues as static modification. The false discovery rate (FDR) for peptide-, protein-, and PSM-level analyses was set to 0.01. All tolerances were set to dynamic for pulsar searches. The XIC extraction for the MS1 and MS2 mass tolerance strategy was set to relative 7 ppm for the timsTOF Pro 2 raw data analysis. The precursor Q-value limit was set to 0.01. Files were normalized using the local normalization strategy on the human proteome. Protein inference was resolved using the IDPicker algorithm. A detailed parameter table is provided in the supplemental table 4.

### Raw data analysis – Utrecht site

Raw data analysis was conducted using DIA-NN 1.9 (Demichev *et al*, 2020). Carbamidomethylation was set as fixed modification while N-terminus cleavage and oxidation were set as variable modifications. Up to two miss-cleavages and one variable modification were allowed. MS1 and MS2 accuracies were set at 10 and 20 ppm, respectively. All other parameters were kept default. A detailed parameter table is provided in the supplemental table 5. The raw data was searched against an in-house blood protein database. This database includes the SwissProt sequences of the 2,445 proteins that are found in blood quantified, by either ELISA or MS, in blood according to the Human Protein Atlas (acquired November 13 2023), to which have been added the top 5% most frequent single amino-acid variations found in the NextProt (Zahn-Zabal *et al*, 2019) database (acquired May 2024). In addition, immunoglobulin sequences have been manually added for a total of 1178 canonical sequences and 1267 mutation sequences.

Proteotypic precursors and proteins groups were filtered at 1% for Q.Value, Lib.Q.Value and Lib.PG.Q.Value. The reminding precursors were then aggregated in protein intensities, based on the Normalized.Precursor intensities, using the MaxLFQ function of the DIAgui R-package (Gerault *et al*, 2024).

### Data Analysis – Utrecht site

Protein concentrations were extrapolated from the protein intensities through a pseudo-calibration approach. Briefly, housekeeping proteins with less than 30% RSD observed through two different cohorts (Gaither *et al*, 2020; Bizzarri *et al*, 2025) were used as reference proteins with their averaged absolute concentrations measured in health donors and considered as theoretical concentrations. A double log_10_ calibration curve was created reporting the theoretical concentration of these house-keeping proteins versus their intensity measured in a pooled serum (HSER-CUSTOM, amsbio) constituted of 400 healthy donors and prepared 4-6 times per sample plate (58 replicates in total). The MaxLFQ intensities were then converted into molar concentration before being converted into mass concentrations using the molar mass of the canonical protein.

### Data analysis - Greifswald site

The Spectronaut report was used as input for SpectroPipeR v0.43 (Michalik *et al*, 2025) to obtain batch-adjusted peptide intensities using the ComBat approach, based on the empirical Bayes methodology (Johnson *et al*, 2007) in the sva package (Leek *et al*, 2012), excluding methionine-oxidized peptides, as well as for the generation of MaxLFQ intensities. The fasta file was used to generate the number of theoretical peptides per protein using the chainsaw tool from the ProteoWizard package (Kessner *et al*, 2008). The sum of the peptide intensities was devided by the number of theoretical peptides to obtain the iBAQ intensities. To identify problematic plasma samples and enables effective quality control the sample quality the PlasmaProteomeProfiling (Geyer *et al*, 2016) gene set list was downloaded from github (https://github.com/MannLabs/Quality-Control-of-the-Plasma-Proteome/blob/master/data/Marker%20List.xlsx). Using this marker list protein proportions per sample were computed and outliers were flagged that exceeded three standard deviations above the harmonic mean (supplemental figure 1). Keratins and protein entries containing variable immunoglobulin regions were removed. The log_10_-scaled protein intensities were utilized to determine the absolute concentrations, employing the log_10_-scaled vegetative plasma protein concentrations proposed by Gaither et al., (Gaither *et al*, 2020) in a linear model. The model was calculated for various quantification methods (iBAQ ∼ g/L; iBAQ ∼ nM; MaxLFQ ∼ g/L; MaxLFQ ∼nM). In order to adjust protein intensities from the 2 sites the global average of the concentration per protein was calculated per site. A ratio was formed between sites yielding in a normalization factor for the Greifswald site. The protein-wise adjustment was subsequently performed by multiplying the adjustment factors with the nanomolar quantities from the Greifswald site. The downstream data analysis and visualization of the data was performed using R (version 4.4.1) (Team, 2014) with various packages (supplemental table 6).

### Multi-classification model

Plasma proteome concentration data were obtained from the Utrecht subset excluding the outlier sample (Trp7_27). Missing values were replaced by random draws from the lower 5th percentile of the observed concentration distribution. To assess the ability of protein profiles to distinguish between individual donors, a multi-class classification approach was employed using a support vector machine (SVM) with a radial basis function (RBF) kernel, implemented via the kernlab package (Karatzoglou *et al*, 2004) in R. For each iteration, a leave-one-sample-per-donor-out strategy was employed: one sample per donor was randomly selected as the test set, while the remaining samples constituted the training set.

Prior to model training, all predictor variables were standardized to have zero mean and unit variance. The modeling pipeline was constructed using the tidymodels (Max Kuhn and Hadley Wickham, 2020) framework, with a fixed cost hyperparameter (C = 1). Probabilistic class predictions were obtained for each test sample, and the predicted class labels were derived from the maximum probability class. Model performance was evaluated across 1,000 independent iterations. For each run, accuracy, sensitivity, and specificity were computed. Sensitivity and specificity were macro-averaged across all classes to ensure equal weighting of each donor, regardless of sample size imbalance.

## Results

Over the past decades, advancements in mass spectrometry have transformed the landscape of plasma proteomics. These technological developments have increased both sample throughput and proteomics depth, paving the way for the analysis of much larger cohorts in plasma proteomics biomarker research. Despite these innovations, many biomarker studies continue to rely on cross-sectional designs, typically measuring only a single time-point per subject. This approach overlooks the potential valuable insights provided by longitudinal sampling, which can reveal donor-specific features and temporal variations critical for understanding dynamic biological processes.

To demonstrate the potential of longitudinal sampling, the TIMES cohort (<u>T</u>racking Individuals <u>M</u>onthly for <u>E</u>valuating <u>S</u>tability) was established (Figure 1A, supplemental table 1). This cohort of 51 participants consisted of 32 females (62.7%) and 19 males (37.3%), was monitored at monthly intervals over a 12-month period. This high-frequency longitudinal sampling of blood enabled assessment of both intra- and inter-individual variability, as well as the temporal stability of candidate plasma biomarkers across the study duration. The supposedly healthy participants had a mean age of 34.33 years (SD = 9.96; range: 19–56 years) and a mean body mass index (BMI) of 24.58 kg/m^2^ (SD = 4.36; range: 18.4–39.8 kg/m^2^) (supplemental table 1).

**Figure 1.**
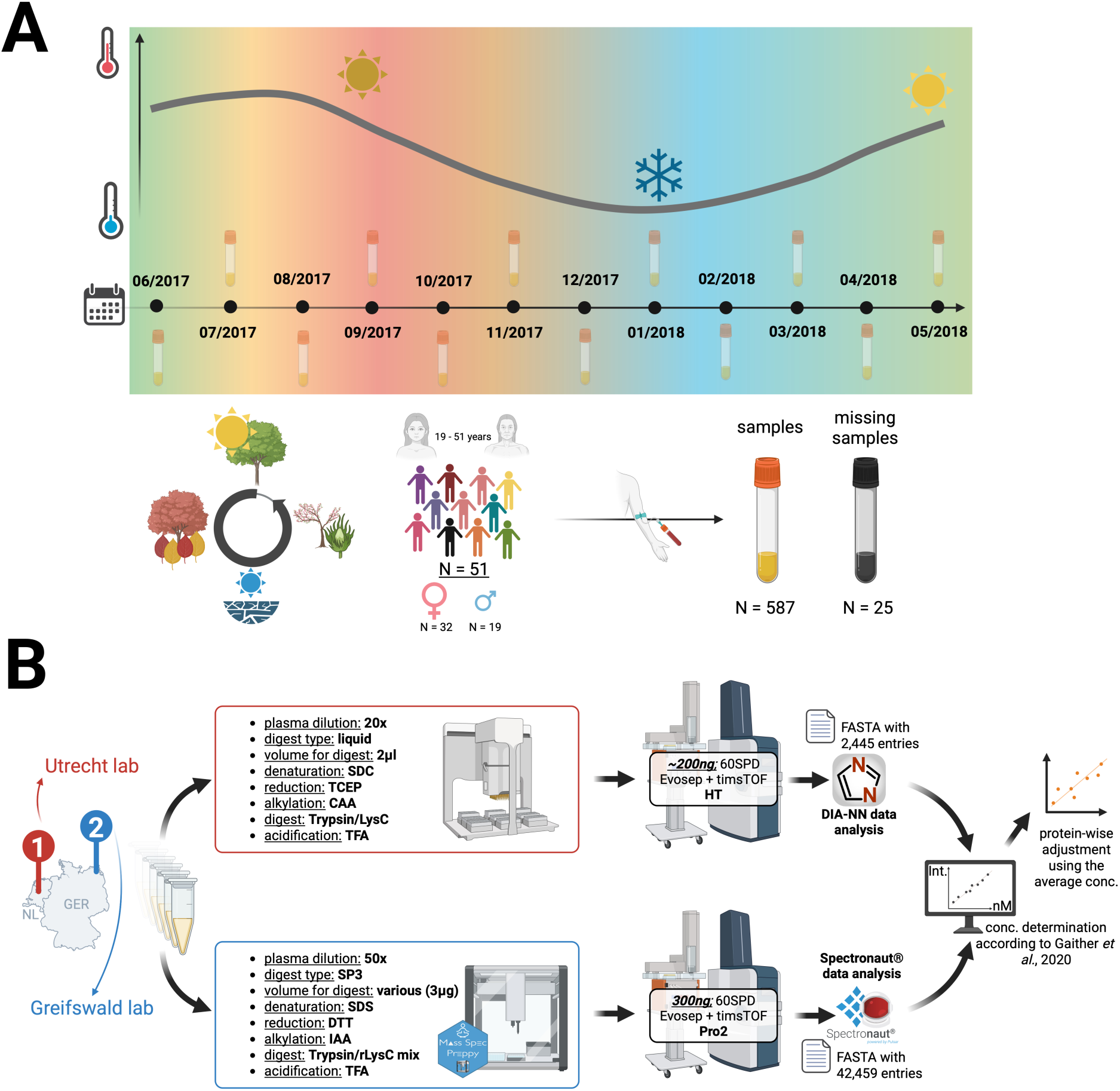
study workflows. Schematic representation of the TIMES Cohort (A) and plasma proteome analysis workflow parameters for each laboratory site (B).

To enable inter-laboratory comparisons, ∼600 plasma samples from the TIMES cohort were analyzed using two independent mass spectrometry workflows at distinct laboratory sites (Figure 1B). This design reflects the challenges of cross-site data comparison in clinical proteomics, where methodological differences including digestion protocols – liquid phase versus bead-based, protein normalized samples versus fixed volume digestion, the sample injection amount load, and mass spectrometry platforms employed. Beyond these experimental factors, data processing introduced further complexity; choices regarding raw data analysis software (e.g. DIA-NN (Demichev *et al*, 2020) versus Spectronaut^®^ (Bruderer *et al*, 2015)), protein database sizes and constructions, and protein quantification methods such as e.g. intensity-based absolute quantification (iBAQ) (Schwanhäusser *et al*, 2011) versus MaxLFQ (Cox *et al*, 2014) were also made independently by each site. Collectively, these layers of methodological variability highlight the inherent challenges in harmonizing proteomic datasets generated across diverse experimental and analytical frameworks.

In this study, the Utrecht site employed a dedicated compact blood-specific protein database comprised of just 2,445 entries in order to avoid false discoveries and decrease search time. This database was built from expected serum proteins labeled as “quantified in plasma” in Human Protein Atlas (Uhlén *et al*, 2015) to which the top 5% most frequent single amino acid mutations of these proteins as reported within the NextProt database were added as new entry with identical gene for compatibility and processing with DIA-NN. In parallel, the Greifswald site utilized a substantially larger human protein database containing 42,459 entries (canonical sequences plus isoforms) in combination with Spectronaut^®^. This also illustrates the diversity of laboratory-specific workflows for plasma proteomics worldwide and, in principle, would be expected to produce comparable results when analyzing identical samples.

Understanding the impact of specific methodological decisions on data quality is therefore critical for optimizing proteomics analysis workflows. To convert relative quantitative data into more clinical-relevant absolute concentration values, absolute quantification data for 37 housekeeping plasma proteins as reported by Gaither *et al*. (Gaither *et al*, 2020) were utilized. These proteins were selected based on the following criteria: a coefficient of variation (CV) less than 0.3 in the Gaither *et al*. dataset (Gaither *et al*, 2020), a CV below 0.3 across the X-omics cohort (n = 150) (Bizzarri *et al*, 2025), and suppression of proteins involved in coagulation pathways to ensure consistency and reliability for both plasma and serum samples. Using these blood plasma housekeeping proteins as reference of known absolute concentration, a log_10_ linear regression was built based on a pooled serum sample of 400 healthy donors at Utrecht Site, whereas Greifswald site independently built the regression individually for every donor. Linear modeling of absolute protein concentrations (either g/L or nM) against relative protein intensities (iBAQ or MaxLFQ) across different acquisition sites demonstrated that iBAQ intensities yielded superior performance regarding the linear model performance in the timsTOF Pro2 dataset, whereas MaxLFQ intensities were more effective in the timsTOF HT dataset (supplemental figure 2). These findings highlight the critical impact of protein quantification and aggregation strategies on the accuracy of absolute concentration estimations. Subsequent analyses were conducted using nanomolar concentration values in combination with iBAQ intensities for the Greifswald site (timsTOF Pro2 dataset) and MaxLFQ intensities for the Utrecht site (timsTOF HT dataset). These combinations were selected based on higher R^2^ values observed in the respective linear models, indicating superior linearity and model performance. Slight differences in protein quantification per sample, which might arise from a combination of workflow-specific parameters, reference sample used for the generation of the calibration line, injection amounts and the dynamic range of the mass spectrometer, resulted in variations for each protein depending on its abundance and detectability within the particular workflow employed. To mitigate these discrepancies, a straightforward approach of averaging protein concentrations per protein across all samples within a single site and building a ratio for each protein between sites yielding normalization factors (Figure 2), which were used to facilitate the adjusting of the data for the comparison of samples across different sites and devices. The majority of high-to mid-abundance proteins exhibit strong concordance, with only minor deviations from a log_2_-ratio of zero, whereas low-abundance proteins demonstrate substantially greater variation in cross–laboratory site ratios. Specifically, analysis across quartiles of averaged median concentration (Q1: lowest to Q4: highest) revealed that only 19.5% of proteins in Q1, 31.7% in Q2, 65.0% in Q3, and 68.3% in Q4 exhibited a cross–laboratory, protein-wise absolute log_2_-ratio magnitude below 1 before adjustment.

**Figure 2.**
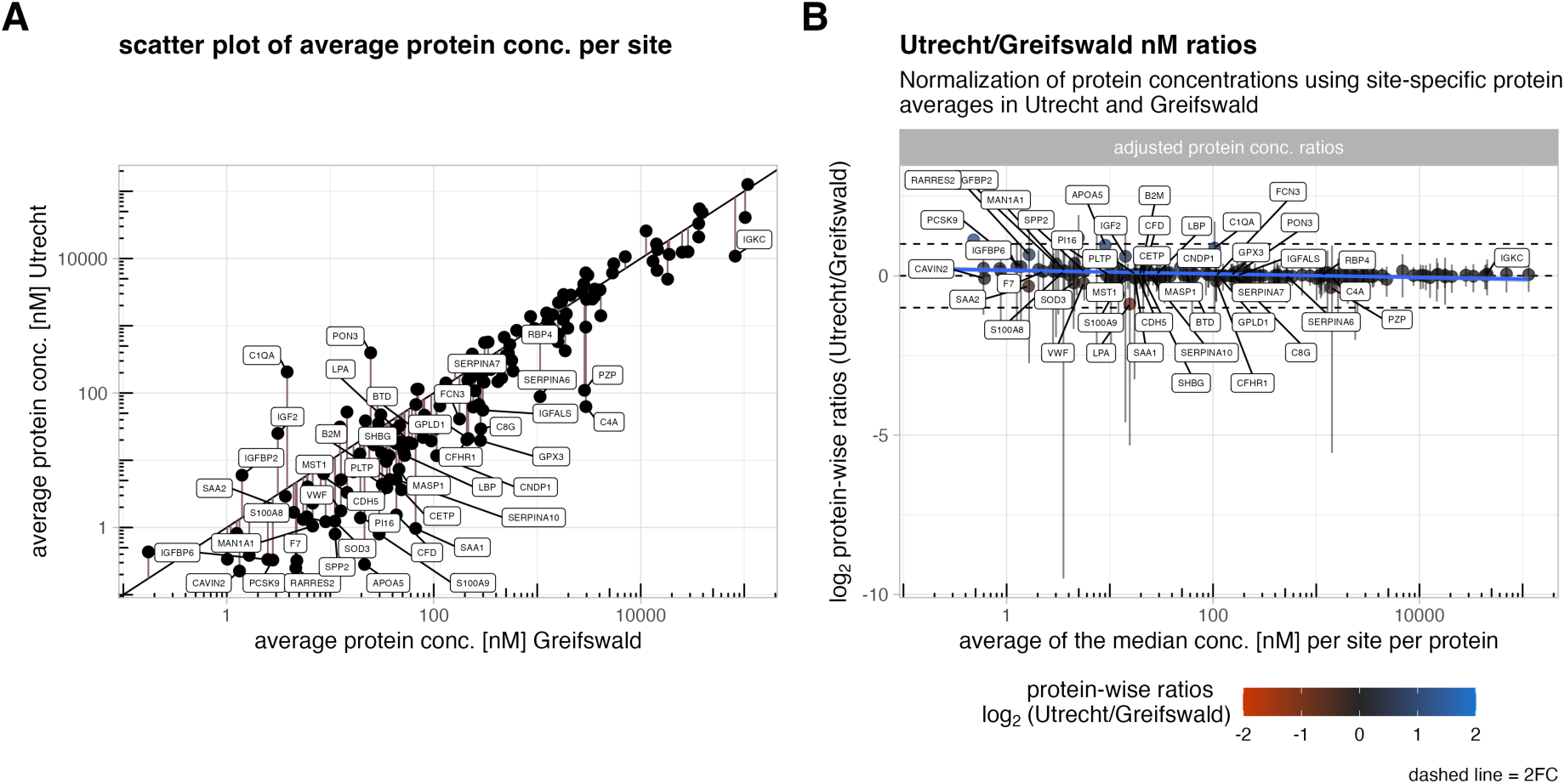
Adjustment of site-specific protein concentrations between Greifswald and Utrecht. Panel A shows a scatter plot comparing average protein concentrations across sites, with vertical pink lines indicating the magnitude and direction of normalization based on adjustment factors. Labels highlight proteins with 4-fold site to site variation. Panel B visualizes log₂ protein-wise concentration ratios after adjustment between Utrecht and Greifswald across protein types, with dashed lines marking 2-fold change thresholds and a fitted linear model illustrating trend behavior. Labels highlight proteins with 4-fold site to site variation.

Overall, after filtering out contaminants and hits containing the variable immunoglobulin regions, 274 proteins at the Greifswald site and 188 proteins at the Utrecht site, could be quantified, with an overlap of 163 proteins. The variations in numbers of proteins covered at both sites reflect the combination of differences in injection amount, sensitivity of MS device, FASTA database composition, and raw data analysis tools.

This stringent filtering led to a rather small subset of genuine plasma proteins, but it still spanned five orders in dynamic range and gave us a large enough dataset to make a sound comparison between the labs for quantitative robustness and reliability. Comparison of protein quantifications between the two laboratory sites on a sample-by-sample basis revealed a strong correlation. Direct pairwise analysis showed that the mean Spearman correlation coefficient across samples was 0.987, while the mean Pearson correlation coefficient was 0.988, highlighting both the consistency of relative protein ranking and the linear agreement in absolute protein abundances between sites (Figure 3).

**Figure 3.**
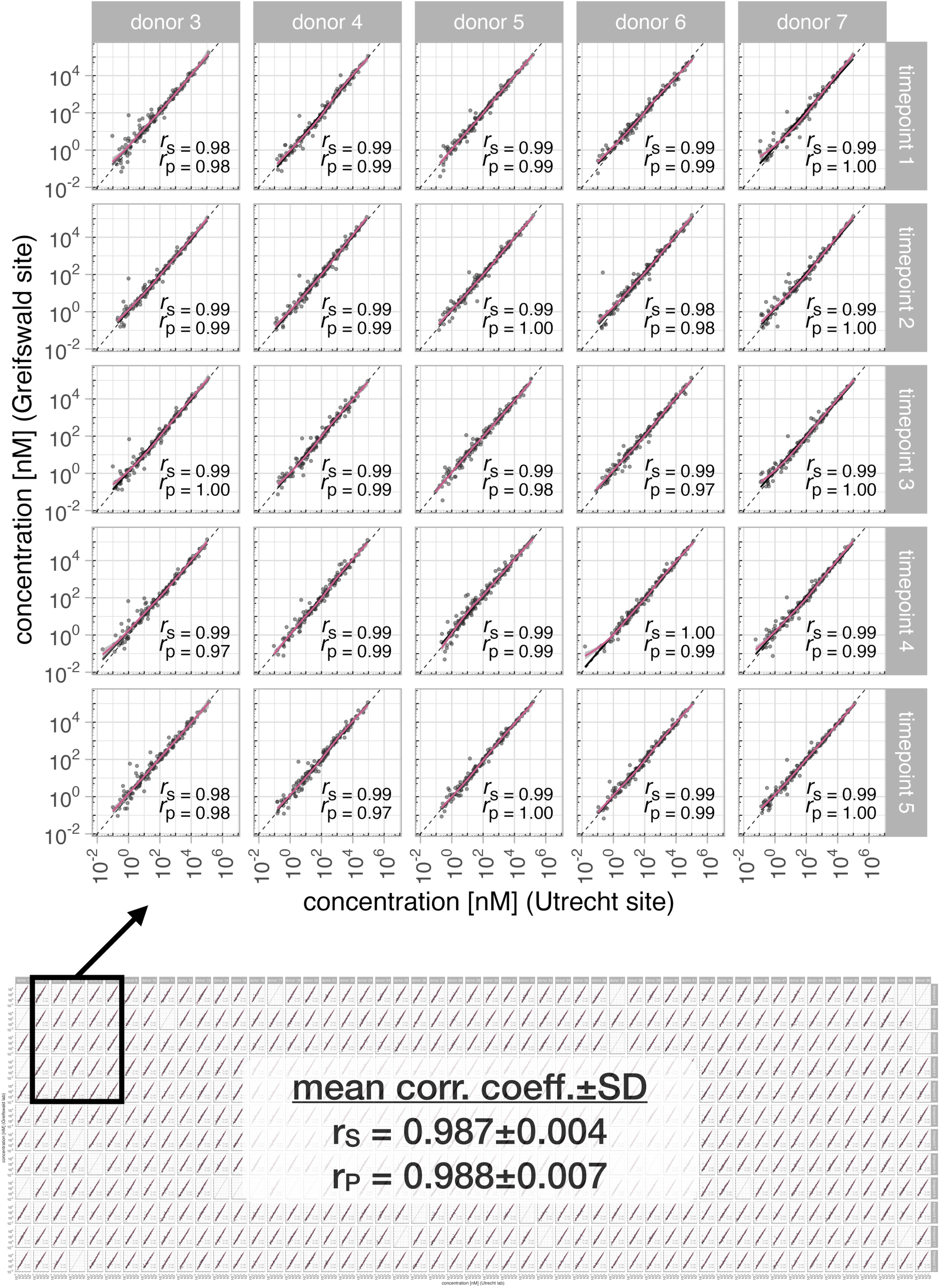
Correlation analysis across lab sites per sample. The overview (lower part) and zoom-in (upper part) panel presents a scatterplot depicting the per-sample measurements between the two lab sites, with Utrecht concentration values on the x-axis and Greifswald concentration values on the y-axis. Spearman and Pearson correlation coefficients (rounded to two decimal places) are shown as rs and rp, respectively.

As we demonstrated a very strong correlation between the two datasets, we opted for further analysis exclusively with the Utrecht site dataset, owing to its superior quantification performance attributed to the specific mass spectrometry device employed. As a reminder, for each donor we recorded 12 plasma proteomes originating from the monthly longitudinal sampling. Uniform Manifold Approximation and Projection (UMAP) analysis of the dataset (Figure 4A) revealed a clear separation between all donors, alongside a consistent protein abundance pattern across monthly sampling for each donor. UMAP is an unsupervised dimensional reduction method that preserves the neighborhood structure of the original high-dimensional data, enabling the visualization of complex relationships in two dimensions. In this context, points that cluster closely in the projection represent samples with highly similar protein abundance profiles across all measured variables prior to dimensional reduction. Accordingly, samples positioned in close proximity within the UMAP plot can be interpreted as sharing comparable protein abundance patterns. In other words, this analysis hints at substantially more inter-donor variation than intra-donor variation. In this context, the longitudinal sampling design of the cohort, combined with the clear donor-specific clustering observed in the UMAP projection, enabled the detection of a potential sample duplication event possibly arising during the aliquoting process.

**Figure 4.**
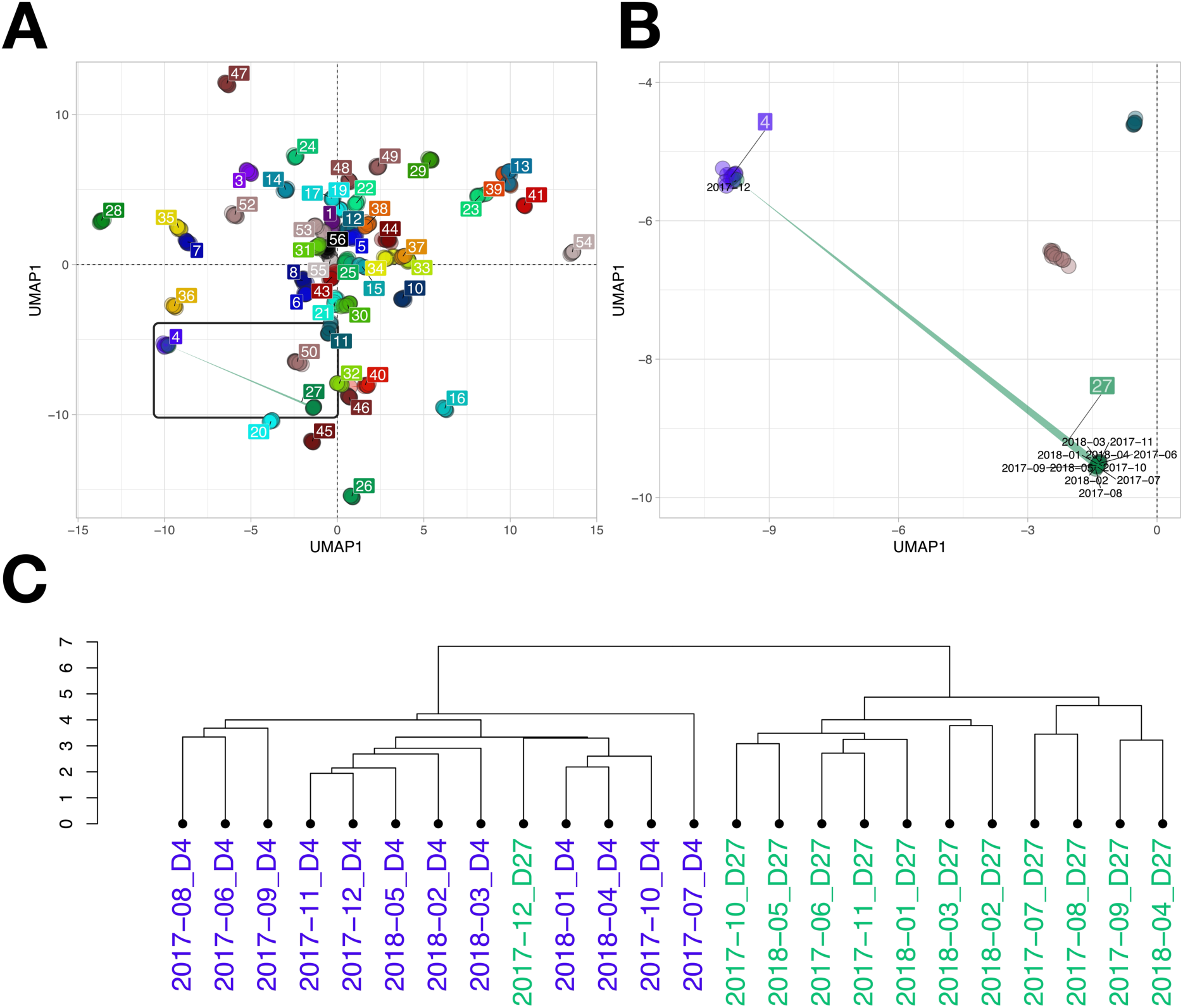
Donor specific protein profiles. Panel A presents a global UMAP projection of plasma protein concentrations, encompassing all donors and their respective monthly samples within the TIMES cohort. Each dot symbolizes a monthly sample, uniquely colored to represent the individual donor. Panel B provides a magnified view of the region highlighted in Panel A, revealing that a sample labeled as donor 27 is embedded within the cluster of donor 4, raising the possibility of sample mislabeling or duplication. Panel C presents hierarchical clustering of protein concentration profiles for both donors, computed using Euclidean distance and complete linkage, further supporting the observed proximity in the reduced dimensional space.

Notably, the December sample of donor 27 was positioned within the point cloud corresponding to donor 4, indicating a protein abundance profile that was indistinguishable from that donor’s longitudinal samples (Figure 4B, C). Given the otherwise robust separation of donors across all time points, this overlap was deemed highly unlikely to occur by chance and was interpreted as evidence of a sample mislabeling or duplication. To avoid introducing artefacts into downstream analyses, this outlier sample was excluded from all subsequent analyses.

To further validate the observation that each donor exhibits a unique and reproducible intra-donor protein profile, enabling donor identification based on the quantitative data, we employed a multi-class SVM classification model (supplemental figure 3), which is a robust methodology for evaluating the ability to distinguish individuals and assessing the consistency of the underlying protein profiles. Across 1,000 iterations, the model achieved a median classification accuracy of 0.98 (Q_1; 25%_ = 0.961 & Q_3; 75%_ = 1). This indicates that, when one sample per donor is withheld and blinded from the training set, the model correctly assigns the sample to the corresponding donor in median 98% of cases.

Immunoglobulins constitute a dominant class of plasma proteins, accounting for approximately 20% of the total protein mass in human blood. Among these, IgG, IgA, and IgM are the most prevalent, with IgG typically representing the most abundant subclass. Human IgG is further subdivided into four structurally related subclasses — IgG1, IgG2, IgG3, and IgG4 — each with distinct sequence motifs, structural properties, and immunological functions. The subclass nomenclature of IgG has been traditionally based on their relative mean concentrations in circulation. Within each subclass exists a heterogeneous repertoire of immunoglobulin clones, ranging from thousands to millions of unique variants, differing subtly in their variable region sequences (Bondt *et al*, 2021). This molecular diversity poses significant analytical challenges for plasma proteomic studies, particularly in database curation and peptide identification workflows. Consequently, the immunoglobulin data are frequently excluded or actively depleted in most reported plasma proteome studies to date, likely to reduce complexity and enhance detection of lower-abundance proteins.

In contrast, our analysis retained the immunoglobulin pool, allowing for quantitation and comparative assessment of these proteins across donors. This approach provides novel insights into the inter-individual variance and underscores the biological relevance of immunoglobulin dynamics in longitudinal plasma proteomics. Despite notable sequence homology among immunoglobulin classes, each exhibits unique peptide sequences, particularly within their Fc domains. These unique regions allow confident identification and semi-quantitative analysis using tryptic peptide profiling. Utilizing peptides unique to each subclass that correspond to the constant heavy chains of IgG1 through IgG4, we were able to estimate the total IgG concentration by assessing the concentration of each subclass and allowed to monitor the temporal consistency of these concentrations. As illustrated in Figure 5, the mass spectrometry-derived concentration estimates showed a strong concordance with established plasma reference ranges for IgG (6.20–15.10 g/L). Across the cohort, the average total IgG concentration across donors was 9.75 g/L, with a standard deviation of 2.64 g/L (27%), and ranged from 3.87 to 16.24 g/L. Strikingly, while inter-donor variability remained substantial (SD over mean total IgG per donor = 2.64 g/L, CV = 0.27, IQR_95%–5%_ = 7.64 g/L), intra-donor fluctuations were minimal (mean SD = 1.11 g/L, mean CV = 0.115, IQR_95%–5%_ = 3.04 g/L), suggesting strong longitudinal stability within individuals over the one-year sampling period.

**Figure 5.**
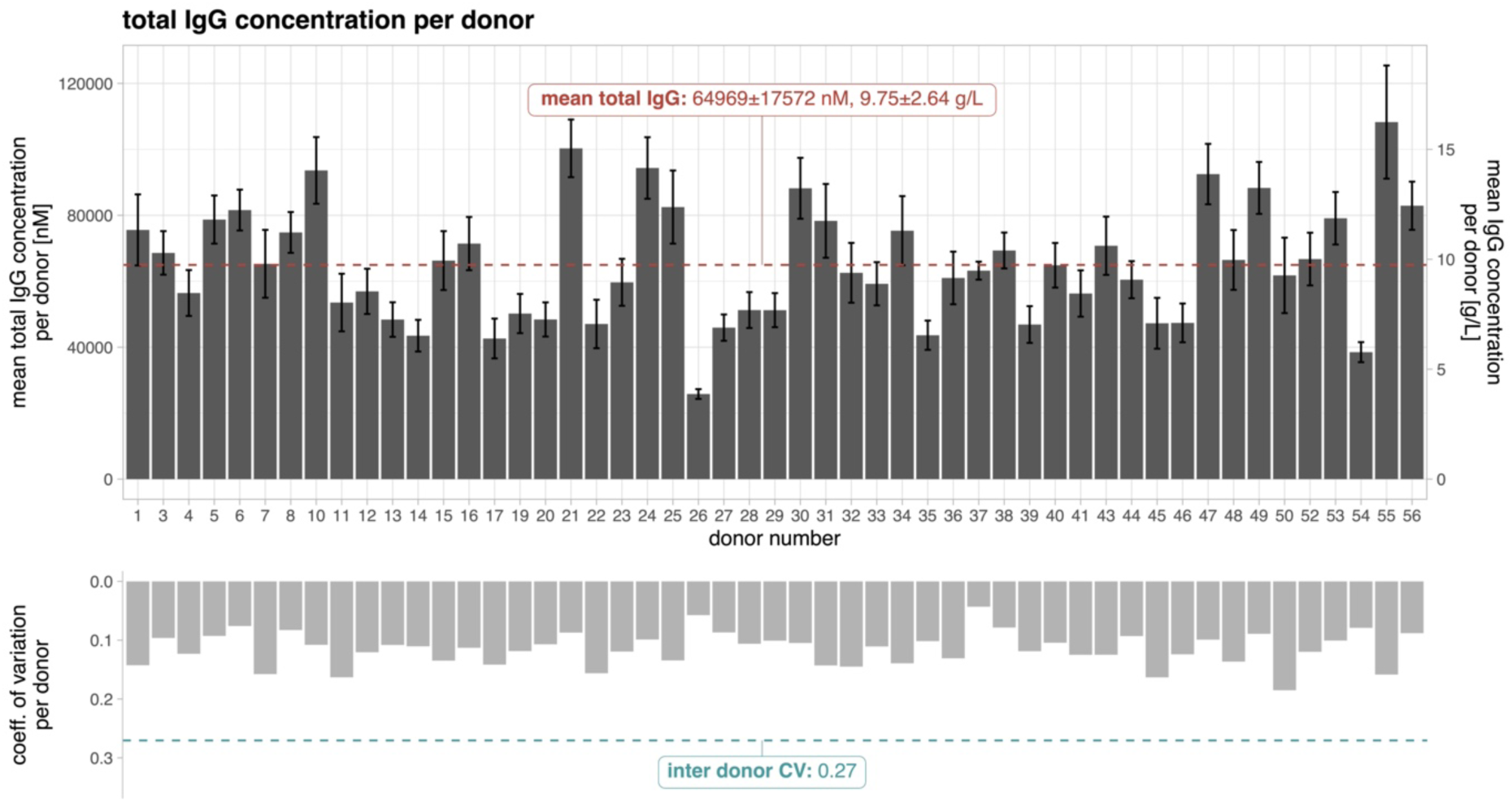
Total IgG profiles reveal pronounced inter-donor variability. Total IgG concentrations were derived by summing all measured IgG sub-classes (IgG1–IgG4) for each donor at every sampling time point, yielding cumulative IgG levels in nanomolar (nM) units. The resulting bar plot depicts the mean and standard deviation of these total IgG concentrations across time points for each individual. Corresponding values in grams per liter (g/L) were computed from nM concentrations using molecular weight–based conversion. The inter-donor coefficient of variation (CV) is depicted as a dashed turquoise line, and its exact value is provided in the label. Similar data presentations are shown in supplemental figure 5-12, but then categorized by immunoglobulin (sub)classes.

Similarly, the four IgG sub-classes showed strong inter-individual variability, but only minor intra-donor fluctuations (Figure 6, supplemental figure 4-13), reinforcing the notion that immunoglobulin levels are tightly regulated under homeostatic conditions. This pattern was particularly pronounced among immunoglobulins, most notably IgG3, IgG4, IgM, IgA2, and IgD. Besides, our data revealed a distributional pattern of CD5L that closely mirrored that of IgM, consistent with prior evidence identifying CD5L as an integral and ubiquitous constituent of plasma IgM complexes (Oskam *et al*, 2023; Wang *et al*, 2024). The observed stability likely reflects the health status of all donors enrolled in the study.

**Figure 6.**
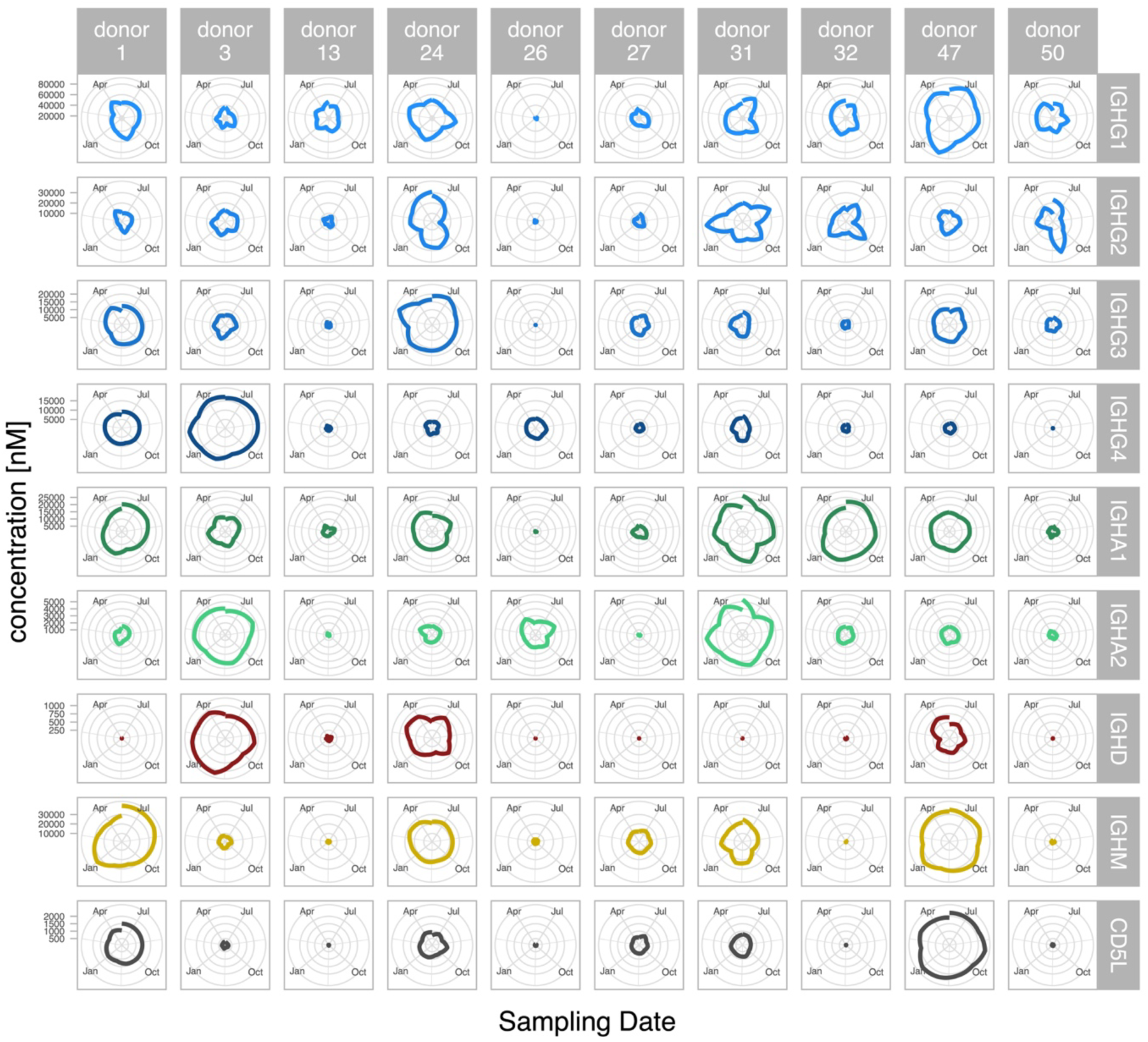
Immunoglobulin sub-classes and CD5L intra- and inter donor variability over time. Nanomolar (nM) concentrations of immunoglobulin sub-classes and CD5L are visualized per donor using radar plots. In this representation, a smaller circular diameter indicates a lower relative concentration compared to other donors. The closer the plotted line approximates a circle, the more stable the concentration remains over the 12-month observation period. Of note, as depicted in this figure donor 26 is characterized by overall low levels of nearly all classes of monitored Ig’s (with IgA2 being the exception). These radarplot representations also illustrate nicely the huge inter donor variation in specific Ig levels, to illustrate for IgD, donors 3, 24 and 47 have relatively high levels of IgD, whereas in the other donors this Ig class is barely detectable. The patterns of Ig (sub) class levels in plasma are highly individual. There is little correlation between them except for IgM and the associated protein CD5L.

Since immunoglobulins are frequently excluded from contemporary proteomics studies, we benchmarked our data against results from an ELISA-based analysis of 600 healthy donors (Gonzalez-Quintela *et al*, 2008). That study reported an average total IgG concentration of 11.1 g/L with a standard deviation of 2.5 g/L (25%), ranging from 7.0 to 17.4 g/L, whereas another study reported a range of 6.20–15.10 g/L (Khan *et al*, 2021). Literature-based adult Ig subclass distributions (Madassery *et al*, 1988; Schauer *et al*, 2003) suggest approximate concentrations of IgG1: 8.3 mg/mL (∼63.6% of total IgG), IgG2: 3.2 mg/mL (∼24.5% of total IgG), IgG3: 0.74 mg/mL (∼5.7% of total IgG), and IgG4: 0.82 mg/mL (∼6.3% of total IgG). In comparison, our mass spectrometry-based quantification yields an average IgG1 at 6.12 mg/mL (62.77% of total IgG), IgG2 at 1.86 mg/mL (19.08%), IgG3 at 0.91 mg/mL (9.33%), and IgG4 at 0.85 mg/mL (8.72%) (supplemental figure 14).

A major observation in our study is that most proteins display very little change in plasma concentration when monitored at regular intervals over the time span of a year. There are however several notable exceptions, exemplified by well-known inflammation markers, such as C-reactive protein (CRP), SAA1/SAA2, and S100A8 and S100A9. For instance, CRP concentrations were generally quite low but exhibited pronounced temporal increases, albeit mostly just in a few donors (Figure 7) resulting in substantial averaged differences between donors. The average total CRP concentration across donors was 19.25 nM, with a standard deviation of 34 nM (176%), and ranged from 0.28 to 209.16 nM (∼750x increase). The annual intra-donor fluctuations were minimal from 0.10-0.39 nM (donor 10) to maximal 0.16-721.43 nM (donor 48) (Figure 7). Of interest, a few donors (13, 26, 34, 37, 49 and 54) revealed a relatively high basal CRP concentration of ∼50 nM, being that high across all longitudinal sampling points. In contrast, donors 21, 33, 48 and 56 had across the year rather low levels of CRP (< 5 nM), albeit that at discrete sampling points the CRP concentrations temporally increased to some of the highest levels measured (i.e. > 200 nM). These latter instances we hypothesize could be linked to short-term events of acute inflammation. Additionally, the inter- and intra-variation patterns observed for CRP across all donors and time points, were consistently observed for the functionally related inflammation markers SAA1, SAA2, and S100A8 and S100A9 (supplemental figure 15-18). The consistency in these episodic peaks observed across all the inflammation markers, strengthens that these observations point at acute phase inflammation events.

**Figure 7.**
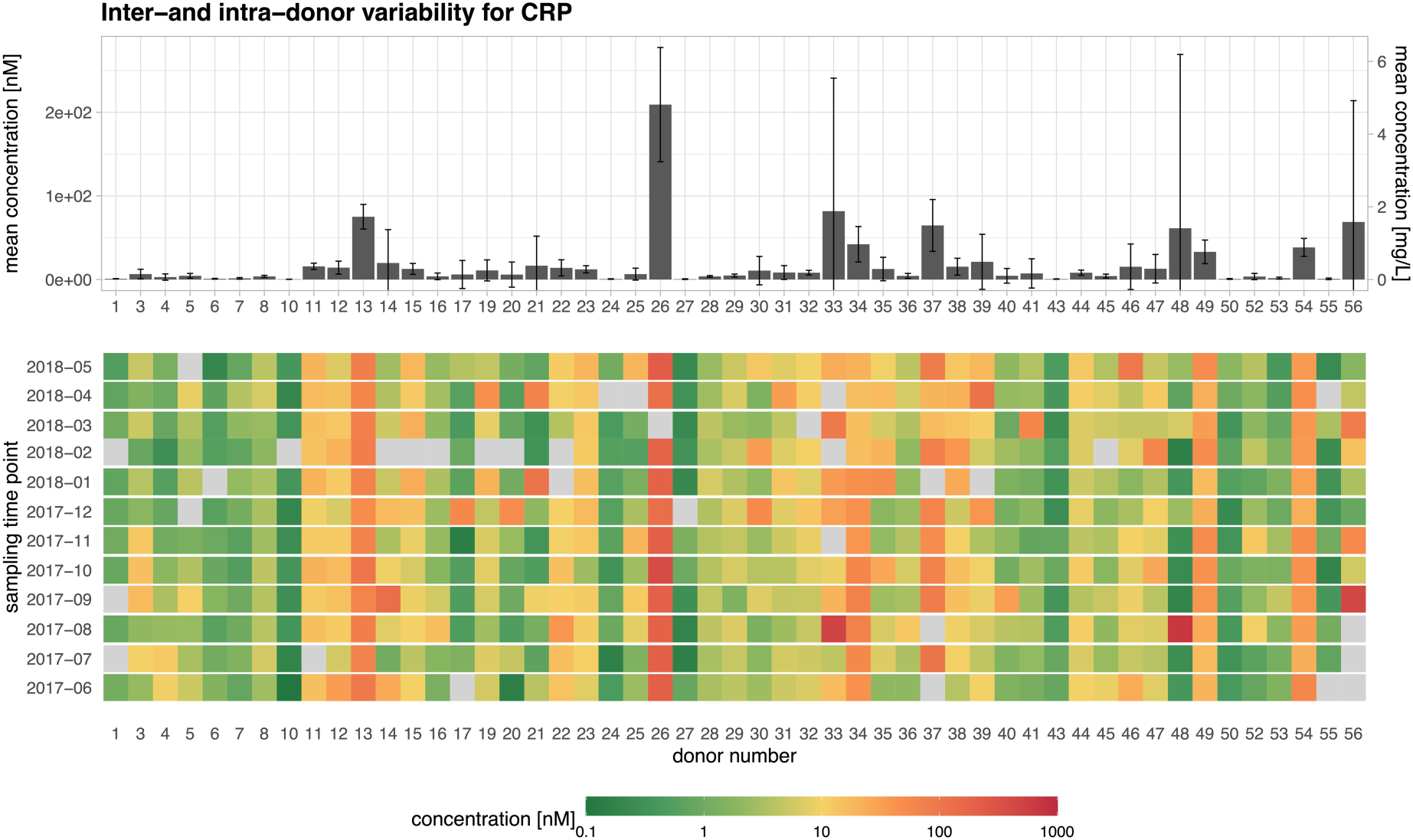
Intra- and inter-donor variability of CRP. CRP mean concentration shown in barplot (top panel). The tile plot below (lower panel) depicts the temporal profiles of CRP in donors, whereby donor 13, 26, 34, 37, 49 and 54 display continuous basal elevated levels of CRP (compared to the average) and donors 21, 33, 48 and 56 show across the year relative low levels of CRP, with temporal bursts of concentration increases at single sampling points.

## Discussion

### Cross-site reproducibility and platform correlation

A major finding, extracted from our collaborative work is that despite substantial differences in sample digestion protocols, injection amounts, instrument dynamic ranges (timsTOF Pro2 vs. timsTOF HT), and data processing workflows, exceptional strong correlation was observed between the datasets acquired at the two sites, even with substantially differing protein identification counts. This correlation has potential for further improvement in the timsTOF HT dataset. Vitko *et al*. (Vitko *et al*, 2024) directly compared the timsTOF Pro2 and timsTOF HT under varying loading amounts and LC gradient conditions. In neat plasma analyses, the timsTOF HT preserved peptide identification rates across increasing sample loads, whereas the timsTOF Pro2 exhibited marked losses, particularly with short LC gradients at >300 ng relative to 100 ng. Quantitative assessment demonstrated MS2 signal attenuation with increasing load on both platforms, with a more pronounced effect for the timsTOF Pro2. At ≥300 ng, the timsTOF HT yielded narrower signal intensity distributions and lower coefficients of variation (CVs), indicating enhanced reproducibility. These results indicate that a sample loading greater than 200 ng is expected to yield a protein identification count comparable to that obtained with the timsTOF Pro2 at 300 ng using the 60 SPD LC workflow. Despite the different mass spectrometers used in our study in combination with different sample loading amounts and at the same time using the same LC setup and gradient, the workflow also varied regarding sample preparation as one site used SP3 sample digest (Hughes *et al*, 2019) whereas the other used a liquid digest approach. SP3 uses paramagnetic beads for protein capture, allowing removal of detergents and other contaminants through multiple wash steps before on-bead digestion. This approach generally improves peptide recovery from low-input samples, enhances reproducibility, and minimizes sample losses during transfer (Hughes *et al*, 2019). In contrast, conventional liquid digestion (in-solution digestion without bead capture) relies on precipitation or dilution to remove detergents, which can lead to variable recovery and reduced reproducibility, particularly at low protein amounts. Liquid digest methods are simpler in workflow but more susceptible to incomplete detergent removal and peptide loss during cleanup. SP3 and SDC in solution digest methods achieved broadly comparable overall proteome coverage; large overlaps have been seen across various bottom methods generally (Varnavides *et al*, 2022). However, SP3-derived samples displayed distinct patterns regardless of the applied buffer systems, suggesting that magnetic bead–mediated protein pulldown constitutes a major determinant of method-specific protein profiles (Varnavides *et al*, 2022). Altogether, these observations indicate that normalization of the data by protein is required to enable valid comparisons between workflows. Although averaging across proteins and adjusting the mean values could facilitate such comparisons, the inter-laboratory evaluation results strongly support including a universal plasma reference sample or using spike-in standards with each measurement workflow (Cai *et al*, 2025) since cohorts and disease phenotype plasma proteome can strongly vary. This approach would allow adjustment of protein intensities independent of the sample preparation, mass spectrometry device and analysis workflow, enabling comparability across cohorts ensuring standardization and reproducibility in multi-center studies in the future.

### Intra- and inter-individual plasma proteome variability

The longitudinal study design of the TIMES cohort revealed that inter-individual differences in plasma protein quantification largely exceed intra-individual variability over time per protein. Complementary findings from a longitudinal study of healthy Danish adults demonstrated that total IgG concentrations, along with most IgG subclasses, remained remarkably stable within individuals over a period of at least 42 weeks (Rasmussen *et al*, 2021). These results underscore the robustness of individual immunoglobulin profiles and support their potential utility in longitudinal biomarker studies and personalized health monitoring (Rasmussen *et al*, 2021). These findings also suggest that a single measurement of IgG subclass levels is generally sufficient to represent an individual’s immunoglobulin profile, even across seasonal variations. But, although the individual proteome remains relatively stable over time, substantial variability exists between donors. This longitudinal cohort also highlights the donor-dependent baseline difference in some proteins of clinical relevance such as CRP. This inter-individual heterogeneity can obscure true biological signals in clinical studies if not properly accounted for. To mitigate this, it would be essential for clinical proteomic analyses to incorporate donor- or patient-specific control samples and express results as relative ratios. Such normalization strategies help adjust for baseline differences and enhance the sensitivity, accuracy and interpretability of biomarker discovery and disease profiling.

## Conclusion

Our study underscores the potential of longitudinal sampling in plasma proteomics, providing deep insights into both intra- and inter-individual variations over time. By leveraging independent mass spectrometry workflows and robust data analysis pipelines, we highlighted the challenges and solutions in cross-laboratory data harmonization. The observed stability of immunoglobulin subclasses contrasts starkly with the dynamic fluctuations of proteins like CRP, reflecting the complexity of human plasma proteomics and interpretation of the results. These findings stress the importance of longitudinal designs for accurately capturing the dynamic biological landscape, which is critical for biomarker discovery research and clinical applications.

### Limitations of the study

Despite the robust design and comprehensive data generated in this study, several limitations need to be acknowledged. First, the cohort size, although sufficient for detecting intra- and inter-individual variations, may limit the generalizability of the findings to broader populations with different demographic or health profiles. Further longitudinal studies with larger, more diverse cohorts are needed to validate and extend these findings. Secondly, we here focus solely on the ∼200 plasma proteins that are consistently detected and can be robustly quantified over all donors and all time points. It has been shown that with more efforts the detectable plasma proteome can be increased to over 1000 proteins, where the gain comes mostly from lower abundant proteins. However, we did not aim for that as it would substantially increase the number of missing values, and thus also CVs and robustness in quantification.

## Resource availability

Restrictions apply to the availability of data generated or analyzed during this study to preserve patient confidentiality or because they were used under license. Access to the data is permitted under controlled access in accordance with the European Union General Data Protection Regulation (GDPR) at https://transfer.ship-med.uni-greifswald.de/FAIRequest/?lang=en.

## Acknowledgments

Schemes were generated using BioRender. The authors would like to thank Prof. Dr. Thomas Thiele and the members of the blood donation center of the University Medicine Greifswald for their support in sample acquisition. We would also like to thank Ulrike Lissner, Erika Friebe, Susanne Neumeister, Sabine Prettin, Fawaz Alsholui, Hanna Friedrich, Christopher Wirks for supporting the conduction of the study, sample processing, sample archiving or sample preparation. This research was partly funded by the Dutch Research Council NWO, funding the Netherlands Proteomics Centre facilities through the X-omics National Road Map program (project 184.034.019) and also supported by the Bundesministerium für Bildung und Forschung (BMBF) via a grant to the German Center of Cardiovascular Research, partner site Greifswald (project 81Z0400101 granted to UV).

## Author contributions

- SM – conceptualization of the study, assistance in sample preparation, data curation, formal analysis, data visualization, writing – original draft, writing – review & editing
- SKN, assistance in sample preparation, data curation, formal analysis, assistance with data visualization, writing – review & editing
- ND, assistance in sample preparation, data curation, formal analysis, assistance with data visualization, writing – review & editing
- MGS, assistance in sample preparation, data curation, writing – review & editing
- EH, assistance in sample preparation, data curation, writing – review & editing
- VD, assistance in sample preparation, data curation
- SH, conceptualization of the study, assistance in sample preparation, data curation, formal analysis, writing – review & editing
- SW, data curation, formal analysis
- HT, assistance in data analysis, data curation, review and editing,
- BB, conceptualization of the study, writing, review and editing of manuscript, supervision and funding acquisition
- GD, conceptualization of the study, assistance in sample preparation, writing – review & editing
- UV, conceptualization of the study, writing, review and editing of manuscript, supervision and funding acquisition
- AJRH, conceptualization of the study, data curation, data visualization, writing – original draft, writing – review & editing, supervision and funding acquisition

## Supplemental information

The supplemental material contains:

- TIMES cohort - study design table
- timsTOF - diaPASEF settings tables
- Spectronaut® settings table
- DIA-NN settings table
- used R packages table
- Plama Proteome Profiling figure
- R-squared of linear modeling figure
- multi-class sample prediction figure
- immunoglobulin sub-types and CD5L figures
- single plots IgG, IgA subtypes and IgM, IgD and CD5L figures
- IgG subclass distribution figure
- selected inflammation markers figures

## References

Aebersold R & Mann M (2016) Mass-spectrometric exploration of proteome structure and function. Nature 537: 347 355

Álvez MB, Bergström S, Kenrick J, Johansson E, Åberg M, Akyildiz M, Altay O, Sköld H, Antonopoulos K, Apostolakis E, et al (2025) A human pan-disease blood atlas of the circulating proteome. Science: eadx2678

Ameling S, Auwera SV der, Holtfreter S, Wiechert A, Michalik S, Friedrich N, Hammer E, Völzke H, Nauck M, Grabe HJ, et al (2025) Cytokine atlas of the population-based cohort SHIP-TREND-0 – Associations with age, sex, and BMI. Cytokine 189: 156896

Anderson NL (2002) The Human Plasma Proteome: History, Character, and Diagnostic Prospects. Mol Cell Proteomics 1: 845 867

Bizzarri D, Akker EB van den, Reinders MJT, Pool R, Beekman M, Lakenberg N, Drouin N, Stecker KE, Heck AJR, Knol EF, et al (2025) Extreme MetaboHealth scores in three cohort studies associate with plasma protein markers for inflammation and cholesterol transport. Immun Ageing 22: 34

Bondt A, Hoek M, Tamara S, Graaf B de, Peng W, Schulte D, Rijswijck DMH van, Boer MA den, Greisch J-F, Varkila MRJ, et al (2021) Human plasma IgG1 repertoires are simple, unique, and dynamic. Cell Syst 12: 1131–1143.e5

Bruderer R, Bernhardt OM, Gandhi T, Miladinović SM, Cheng L-Y, Messner S, Ehrenberger T, Zanotelli V, Butscheid Y, Escher C, et al (2015) Extending the limits of quantitative proteome profiling with data-independent acquisition and application to acetaminophen treated 3D liver microtissues. Mol Cell Proteomics 14: 1400 1410

Cai X, Geyer PE, Perez-Riverol Y, Omenn GS, Dong L, Winkler R, Ahadi S, Lössl P, Yu X, Chang C, et al (2025) A standardized framework for circulating blood proteomics. Nat Genet: 1–10

Cox J, Hein MY, Luber CA, Paron I, Nagaraj N & Mann M (2014) Accurate proteome-wide label-free quantification by delayed normalization and maximal peptide ratio extraction, termed MaxLFQ. Mol Cell Proteomics 13: 2513 2526

Demichev V, Messner CB, Vernardis SI, Lilley KS & Ralser M (2020) DIA-NN: neural networks and interference correction enable deep proteome coverage in high throughput. Nat Methods 17: 41–44

Dou YN, Grimstein C, Mascaro J & Wang J (2024) Biomarkers for Precision Patient Selection in Cancer Therapy Approvals in the US, from 2011 to 2023. Clin Pharmacol Ther 116: 304–314

Gaither C, Popp R, Mohammed Y & Borchers CH (2020) Determination of the concentration range for 267 proteins from 21 lots of commercial human plasma using highly multiplexed multiple reaction monitoring mass spectrometry. Analyst 145: 3634–3644

Gerault M-A, Camoin L & Granjeaud S (2024) DIAgui: a Shiny application to process the output from DIA-NN. Bioinform Adv 4: vbae001

Geyer PE, Holdt LM, Teupser D & Mann M (2017) Revisiting biomarker discovery by plasma proteomics. Mol Syst Biol 13: 942

Geyer PE, Kulak NA, Pichler G, Holdt LM, Teupser D & Mann M (2016) Plasma Proteome Profiling to Assess Human Health and Disease. Cell Syst 2: 185–195

Gonzalez-Quintela A, Alende R, Gude F, Campos J, Rey J, Meijide LM, Fernandez-Merino C & Vidal C (2008) Serum levels of immunoglobulins (IgG, IgA, IgM) in a general adult population and their relationship with alcohol consumption, smoking and common metabolic abnormalities. Clin Exp Immunol 151: 42–50

Hendricks NG, Bhosale SD, Keoseyan AJ, Ortiz J, Stotland A, Seyedmohammad S, Nguyen CDL, Bui JT, Moradian A, Mockus SM, et al (2024) An Inflection Point in High-Throughput Proteomics with Orbitrap Astral: Analysis of Biofluids, Cells, and Tissues. J Proteome Res 23: 4163–4169

Hughes CS, Moggridge S, Müller T, Sorensen PH, Morin GB & Krijgsveld J (2019) Single-pot, solid-phase-enhanced sample preparation for proteomics experiments. Nat Protoc 14: 68–85

Johnson WE, Li C & Rabinovic A (2007) Adjusting batch effects in microarray expression data using empirical Bayes methods. Biostatistics 8: 118 127

Karatzoglou A, Smola A, Hornik K & Zeileis A (2004) kernlab - An S4 Package for Kernel Methods in R. J Stat Softw 11

Kessner D, Chambers M, Burke R, Agus D & Mallick P (2008) ProteoWizard: open source software for rapid proteomics tools development. Bioinformatics 24: 2534 2536

Khan SR, Chaker L, Ikram MA, Peeters RP, Hagen PM van & Dalm VASH (2021) Determinants and Reference Ranges of Serum Immunoglobulins in Middle-Aged and Elderly Individuals: a Population-Based Study. J Clin Immunol 41: 1902–1914

Leek JT, Johnson WE, Parker HS, Jaffe AE & Storey JD (2012) The sva package for removing batch effects and other unwanted variation in high-throughput experiments. Bioinformatics 28: 882 883

Madassery JV, Kwon OH, Lee SY & Nahm MH (1988) IgG2 subclass deficiency: IgG subclass assays and IgG2 concentrations among 8015 blood donors. Clin Chem 34: 1407– 1413

Max Kuhn and Hadley Wickham (2020) Tidymodels: a collection of packages for modeling and machine learning using tidyverse principles.

Messner CB, Demichev V, Wendisch D, Michalick L, White M, Freiwald A, Textoris-Taube K, Vernardis SI, Egger A-S, Kreidl M, et al (2020) Ultra-High-Throughput Clinical Proteomics Reveals Classifiers of COVID-19 Infection. Cell Syst 11: 11–24.e4

Michalik S, Hammer E, Steil L, Salazar MG, Hentschker C, Surmann K, Busch LM, Sura T & Völker U (2025) SpectroPipeR—a streamlining post Spectronaut® DIA-MS data analysis R package. Bioinformatics: btaf086

Nimmerjahn F, Vidarsson G & Cragg MS (2023) Effect of posttranslational modifications and subclass on IgG activity: from immunity to immunotherapy. Nat Immunol 24: 1244– 1255

Nteak SK, Völlmy F, Lukassen MV, Toorn H van den, Boer MA den, Bondt A, Lans SPA van der, Haas P-J, Zuilen AD van, Rooijakkers SHM, et al (2024) Longitudinal Fluctuations in Protein Concentrations and Higher-Order Structures in the Plasma Proteome of Kidney Failure Patients Subjected to a Kidney Transplant. J Proteome Res 23: 2124–2136

Oskam N, Boer MA den, Lukassen MV, Heer PO, Veth TS, Mierlo G van, Lai S-H, Derksen NIL, Yin V, Streutker M, et al (2023) CD5L is a canonical component of circulatory IgM. Proc Natl Acad Sci 120: e2311265120

Patel P, Jamal Z & Ramphul K (2023) Immunoglobulin (PMID: 30035936 Bookshelf ID: NBK513460) Treasure Island (FL): StatPearls Publishing

Rasmussen KF, Sprogøe U, Nielsen C, Shalom D & Assing K (2021) Time-related variation in IgG subclass concentrations in a group of healthy Danish adults. *Immun*, Inflamm Dis 9: 1009–1015

Reder A, Hentschker C, Steil L, Salazar MG, Hammer E, Dhople VM, Sura T, Lissner U, Wolfgramm H, Dittmar D, et al (2023) MassSpecPreppy—An end-to-end solution for automated protein concentration determination and flexible sample digestion for proteomics applications. PROTEOMICS

Schauer U, Stemberg F, Rieger CH, Borte M, Schubert S, Riedel F, Herz U, Renz H, Wick M, Carr-Smith HD, et al (2003) IgG Subclass Concentrations in Certified Reference Material 470 and Reference Values for Children and Adults Determined with The Binding Site Reagents. Clin Chem 49: 1924–1929

Schwanhäusser B, Busse D, Li N, Dittmar G, Schuchhardt J, Wolf J, Chen W & Selbach M (2011) Global quantification of mammalian gene expression control. Nature 473: 337–342

Team RDC (2014) R: A Language and Environment for Statistical Computing

Uhlén M, Fagerberg L, Hallström BM, Lindskog C, Oksvold P, Mardinoglu A, Sivertsson Å, Kampf C, Sjöstedt E, Asplund A, et al (2015) Tissue-based map of the human proteome. Science 347: 1260419

Varnavides G, Madern M, Anrather D, Hartl N, Reiter W & Hartl M (2022) In Search of a Universal Method: A Comparative Survey of Bottom-Up Proteomics Sample Preparation Methods. J Proteome Res

Vitko D, Chou W-F, Golmaei SN, Lee J-Y, Belthangady C, Blume J, Chan JK, Flores-Campuzano G, Hu Y, Liu M, et al (2024) timsTOF HT Improves Protein Identification and Quantitative Reproducibility for Deep Unbiased Plasma Protein Biomarker Discovery. J Proteome Res

Völlmy F, Toorn H van den, Chiozzi RZ, Zucchetti O, Papi A, Volta CA, Marracino L, Sega FVD, Fortini F, Demichev V, et al (2021) A serum proteome signature to predict mortality in severe COVID-19 patients. Life Sci Alliance 4: e202101099

Wang Y, Su C, Ji C & Xiao J (2024) CD5L associates with IgM via the J chain. Nat Commun 15: 8397

Zahn-Zabal M, Michel P-A, Gateau A, Nikitin F, Schaeffer M, Audot E, Gaudet P, Duek PD, Teixeira D, Rech de Laval V, et al (2019) The neXtProt knowledgebase in 2020: data, tools and usability improvements. Nucleic Acids Res 48: D328–D334

Zhang H, Liu AY, Loriaux P, Wollscheid B, Zhou Y, Watts JD & Aebersold R (2007) Mass Spectrometric Detection of Tissue Proteins in Plasma* S. Mol Cell Proteom 6: 64–71

