## supplemental material for "Immunoglobulin sub-class levels define inter-donor plasma variability: a longitudinal dual-lab study"

**- Supplemental Materials -**

Stephan Michalik[1,2,*], Sofia Kalaidopoulou Nteak[3,4,*], Nicolas Drouin[3,4,*], Manuela Gesell Salazar[1], Elke Hammer[1,2], Vishnu Mukund Dhople[1,2], Silva Holtfreter[5], Stefan Weiss[1,2], Henk Toorn[3,4], Barbara M. Bröker[5], Grażyna Domańska[5], Uwe Völker[1,2,#], Albert J. R. Heck[3,4,#]

[1] Interfaculty Institute of Genetics and Functional Genomics, Department Functional Genomics, University Medicine Greifswald, Greifswald, Germany
[2] German center for cardiovascular research (DZHK), partner site Greifswald, Germany
[3] Biomolecular Mass Spectrometry and Proteomics, Bijvoet Centre for Biomolecular Research and Utrecht Institute for Pharmaceutical Sciences, Utrecht University, Utrecht, Netherlands
[4] Netherlands Proteomics Center, Utrecht, the Netherlands
[5] Institute of Immunology, University Medicine Greifswald, Greifswald, Germany

[*] These authors contributed equally.
[#] These authors contributed equally and are corresponding authors.

### Table of contents

**1 TIMES cohort - study design** **8**

**2 timsTOF - diaPASEF settings** **9**
diaPASEF settings timsTOF Pro 2 - Greifswald site . . . . . . . . . . . . . . . . . . . 9
diaPASEF settings timsTOF HT - Utrecht site . . . . . . . . . . . . . . . . . . . . . . 9

**3 Spectronaut® settings** **11**

**4 DIA-NN settings** **15**

**5 used R packages** **17**

**6 Plama Proteome Profiling** **19**

**7 R-squared of linear modeling** **20**

**8 multi-class sample prediction** **21**

**9 immunoglobulin sub-types and CD5L** **22**

**10 single plots IgG, IgA subtypes and IgM, IgD and CD5L** **23**

**11 IgG subclass distribution** **32**

**12 selected inflammation markers** **33**

**References** **37**

Supplemental Figure 5 Bar plot of IgG1 concentrations - The bar plot illustrates the mean and standard deviation of IgG1 concentrations across time points for each individual. Corresponding values in grams per liter (g/L) were derived from nanomolar (nM) concentrations using molecular weight-based conversion. The mean (dashed line) and standard deviation across donors are indicated in the label box. The barplot below presents the coefficient of variation (CV) across time points per donor. The inter-donor coefficient of variation is depicted as a dashed turquoise line, and its exact value is provided in the label. . . . . . . . . . . . . . . . . . . . . . 23

Supplemental Figure 6 Bar plot of IgG2 concentrations - The bar plot illustrates the mean and standard deviation of IgG2 concentrations across time points for each individual. Corresponding values in grams per liter (g/L) were derived from nanomolar (nM) concentrations using molecular weight-based conversion. The mean (dashed line) and standard deviation across donors are indicated in the label box. The barplot below presents the coefficient of variation (CV) across time points per donor. The inter-donor coefficient of variation is depicted as a dashed turquoise line, and its exact value is provided in the label. . . . . . . . . . . . . . . . . . . . . 24
Supplemental Figure 7 Bar plot of IgG3 concentrations - The bar plot illustrates the mean and standard deviation of IgG3 concentrations across time points for each individual. Corresponding values in grams per liter (g/L) were derived from nanomolar (nM) concentrations using molecular weight-based conversion. The mean (dashed line) and standard deviation across donors are indicated in the label box. The barplot below presents the coefficient of variation (CV) across time points per donor. The inter-donor coefficient of variation is depicted as a dashed turquoise line, and its exact value is provided in the label. . . . . . . . . . . . . . . . . . . . . 25
Supplemental Figure 8 Bar plot of IgG4 concentrations - The bar plot illustrates the mean and standard deviation of IgG4 concentrations across time points for each individual. Corresponding values in grams per liter (g/L) were derived from nanomolar (nM) concentrations using molecular weight-based conversion. The mean (dashed line) and standard deviation across donors are indicated in the label box. The barplot below presents the coefficient of variation (CV) across time points per donor. The inter-donor coefficient of variation is depicted as a dashed turquoise line, and its exact value is provided in the label. . . . . . . . . . . . . . . . . . . . . 26
Supplemental Figure 9 Bar plot of IgA1 concentrations - The bar plot illustrates the mean and standard deviation of IgA1 concentrations across time points for each individual. Corresponding values in grams per liter (g/L) were derived from nanomolar (nM) concentrations using molecular weight-based conversion. The mean (dashed line) and standard deviation across donors are indicated in the label box. The barplot below presents the coefficient of variation (CV) across time points per donor. The inter-donor coefficient of variation is depicted as a dashed turquoise line, and its exact value is provided in the label. . . . . . . . . . . . . . . . . . . . . 27

Supplemental Figure 10 Bar plot of IgA2 concentrations - The bar plot illustrates the mean and standard deviation of IgA2 concentrations across time points for each individual. Corresponding values in grams per liter (g/L) were derived from nanomolar (nM) concentrations using molecular weight-based conversion. The mean (dashed line) and standard deviation across donors are indicated in the label box. The barplot below presents the coefficient of variation (CV) across time points per donor. The inter-donor coefficient of variation is depicted as a dashed turquoise line, and its exact value is provided in the label. . . . . . . . . . . . . . . . . . . . . . 28
Supplemental Figure 11 Bar plot of IgD concentrations - The bar plot illustrates the mean and standard deviation of IgD concentrations across time points for each individual. Corresponding values in grams per liter (g/L) were derived from nanomolar (nM) concentrations using molecular weight-based conversion. The mean (dashed line) and standard deviation across donors are indicated in the label box. The barplot below presents the coefficient of variation (CV) across time points per donor. The inter-donor coefficient of variation is depicted as a dashed turquoise line, and its exact value is provided in the label. . . . . . . . . . . . . . . . . . . . . . 29
Supplemental Figure 12 Bar plot of IgM concentrations - The bar plot illustrates the mean and standard deviation of IgM concentrations across time points for each individual. Corresponding values in grams per liter (g/L) were derived from nanomolar (nM) concentrations using molecular weight-based conversion. The mean (dashed line) and standard deviation across donors are indicated in the label box. The barplot below presents the coefficient of variation (CV) across time points per donor. The inter-donor coefficient of variation is depicted as a dashed turquoise line, and its exact value is provided in the label. . . . . . . . . . . . . . . . . . . . . . 30
Supplemental Figure 13 Bar plot of CD5L concentrations - The bar plot illustrates the mean and standard deviation of CD5L concentrations across time points for each individual. Corresponding values in grams per liter (g/L) were derived from nanomolar (nM) concentrations using molecular weight-based conversion. The mean (dashed line) and standard deviation across donors are indicated in the label box. The barplot below presents the coefficient of variation (CV) across time points per donor. The inter-donor coefficient of variation is depicted as a dashed turquoise line, and its exact value is provided in the label. . . . . . . . . . . . . . . . . . . . . . 31

### 1 TIMES cohort - study design

The TIMES study aimed to investigate the annual variations in the plasma proteome of healthy individuals. Eligibility criteria included being between 18 and 60 years old, providing written informed consent, and not having any acute illness within the past seven days before sampling. Exclusion criteria included a body mass index below 18.5 kg/m$^2$, as well as the presence of blood coagulation disorders, anemia, or similar conditions. Plasma samples were collected from a total of 53 participants (with a mean age of 34.4 $\pm$ 10.0 years) once a month for a year, spanning from June 2017 to May 2018. Due to early loss to follow-up, two subjects were excluded from the data analyses. Consequently, data from the remaining 51 participants were included in the analyses. A table with phenotyping data of donors can be find below (Supplemental Table 1).

**Supplemental Table 1:** TIMES cohort overview

| | Overall (N=51) |
|---|---|
| **Sex** | |
| - female | 32 (62.7%) |
| - male | 19 (37.3%) |
| **Age (years)** | |
| - Mean (SD) | 34.333 (9.955) |
| - Range | 19.000 - 56.000 |
| **weight (kg)** | |
| - Mean (SD) | 74.810 (18.758) |
| - Range | 51.000 - 145.000 |
| **size (cm)** | |
| - Mean (SD) | 173.608 (10.971) |
| - Range | 151.000 - 198.000 |
| **sample count per donor** | |
| - Mean (SD) | 11.392 (0.918) |
| - Range | 9.000 - 12.000 |
| **BMI** | |
| - Mean (SD) | 24.578 (4.364) |
| - Range | 18.400 - 39.800 |

### 2 timsTOF - diaPASEF settings

#### diaPASEF settings timsTOF Pro 2 - Greifswald site

**Supplemental Table 2:** diaPASEF windows timsTOF Pro 2

| cycle ID | 1/K0 start | 1/K0 end | m/z start | m/z end |
|---|---|---|---|---|
| 1 | 0.67 | 0.85 | 330.17 | 415.23 |
| 1 | 0.85 | 1.12 | 688.69 | 728.86 |
| 2 | 0.70 | 0.90 | 413.23 | 485.01 |
| 2 | 0.90 | 1.15 | 726.86 | 768.90 |
| 3 | 0.72 | 0.93 | 483.01 | 528.94 |
| 3 | 0.93 | 1.19 | 766.90 | 813.39 |
| 4 | 0.74 | 0.96 | 526.94 | 561.75 |
| 4 | 0.96 | 1.23 | 811.39 | 861.41 |
| 5 | 0.76 | 0.99 | 559.75 | 595.00 |
| 5 | 0.99 | 1.28 | 859.41 | 923.50 |
| 6 | 0.78 | 1.02 | 593.00 | 625.33 |
| 6 | 1.02 | 1.32 | 921.50 | 1001.24 |
| 7 | 0.79 | 1.05 | 623.33 | 656.11 |
| 7 | 1.05 | 1.39 | 999.24 | 1107.20 |
| 8 | 0.80 | 1.08 | 654.11 | 690.69 |
| 8 | 1.08 | 1.47 | 1105.20 | 1638.48 |

#### diaPASEF settings timsTOF HT - Utrecht site

**Supplemental Table 3:** diaPASEF windows timsTOF HT

| cycle ID | 1/K0 start | 1/K0 end | m/z start | m/z end |
|---|---|---|---|---|
| 1 | 0.600 | 0.832 | 301.84 | 407.88 |
| 1 | 0.832 | 1.600 | 621.80 | 641.85 |

*(continued)*

| cycle ID | 1/K0 start | 1/K0 end | m/z start | m/z end |
|---|---|---|---|---|
| 2 | 0.600 | 0.862 | 407.88 | 436.22 |
| 2 | 0.862 | 1.600 | 641.85 | 662.35 |
| 3 | 0.600 | 0.872 | 436.22 | 458.25 |
| 3 | 0.872 | 1.600 | 662.35 | 683.84 |
| 4 | 0.600 | 0.892 | 458.25 | 478.22 |
| 4 | 0.892 | 1.600 | 683.84 | 707.82 |
| 5 | 0.600 | 0.902 | 478.22 | 496.74 |
| 5 | 0.902 | 1.600 | 707.82 | 732.86 |
| 6 | 0.600 | 0.912 | 496.74 | 514.77 |
| 6 | 0.912 | 1.600 | 732.86 | 761.38 |
| 7 | 0.600 | 0.922 | 514.77 | 531.80 |
| 7 | 0.922 | 1.600 | 761.38 | 792.34 |
| 8 | 0.600 | 0.942 | 531.80 | 549.29 |
| 8 | 0.942 | 1.600 | 792.34 | 827.87 |
| 9 | 0.600 | 0.952 | 549.29 | 566.80 |
| 9 | 0.952 | 1.600 | 827.87 | 869.93 |
| 10 | 0.600 | 0.972 | 566.80 | 584.62 |
| 10 | 0.972 | 1.600 | 869.93 | 924.00 |
| 11 | 0.600 | 1.002 | 584.62 | 602.33 |
| 11 | 1.002 | 1.600 | 924.00 | 1001.48 |
| 12 | 0.600 | 1.052 | 602.33 | 621.80 |
| 12 | 1.052 | 1.600 | 1001.48 | 1199.55 |

### 3 Spectronaut® settings

The raw data analysis at the Greifswald site was conducted using Spectronaut® (version 20.1.250624.92449) in directDIA+ mode. The analysis utilized reviewed uniprot databases, specifically the *Homo sapiens* database, which contains 42,529 canonical and isoform entries. A detailed list of all parameters used is provided in the table below (Supplemental Table 4).

**Supplemental Table 4:** Spectronaut® settings

| Level 1 | Level 2 | Level 3 | Level 4 | Value |
|---|---|---|---|---|
| Calibration | Calibration Mode | | | Automatic |
| Calibration | MZ Extraction Strategy | | | Maximum Intensity |
| Calibration | Used Biognosys' iRT Kit | | | FALSE |
| Calibration | Allow source specific iRT Calibration | | | TRUE |
| Calibration | Precision iRT | Exclude De-amidated Peptides | | TRUE |
| Calibration | Precision iRT | iRT <-> RT Regression Type | | Local (Non-Linear) Regression |
| Calibration | Calibration Carry-Over | | | FALSE |
| Calibration | MS1 Mass Tolerance Strategy | | | System Default |
| Calibration | MS2 Mass Tolerance Strategy | | | System Default |
| Identification | Precursor Qvalue Cutoff | | | 0.01 |
| Identification | Precursor Qvalue Cutoff (Experiment) | | | 0.01 |
| Identification | Precursor PEP Cutoff | | | 0.2 |
| Identification | Protein Qvalue Cutoff (Experiment) | | | 0.01 |
| Identification | Protein Qvalue Cutoff (Run) | | | 0.05 |

*(continued)*

| Level 1 | Level 2 | Level 3 | Level 4 | Value |
|---|---|---|---|---|
| Identification | Protein PEP Cutoff | | | 0.75 |
| Identification | Single Hit Definition | | | By Stripped Sequence |
| Identification | Single Hit Protein Rule | | | Stratified Single Hit Protein FDR |
| Identification | Run-Level Protein Scoring | | | All Observations |
| Identification | Exclude Duplicate Assays | | | TRUE |
| Identification | Exclude Predicted Fragment Scores | | | FALSE |
| Identification | Generate Decoys | Decoy Generation Method | Preferred Fragment Source | NN Predicted Fragments |
| Identification | Generate Decoys | Decoy Limit Strategy | Library Size Fraction | 0.1 |
| Identification | Pvalue Estimator | | | Kernel Density Estimator |
| Protein Inference | Protein Inference Workflow | Inference Algorithm | | IDPicker |
| Quantification | Precursor Filtering | Imputation Strategy | | Use Background Signal |
| Quantification | Precursor Filtering | Multi Channel Qvalue Filter | | Group Qvalue |
| Quantification | Proteotypicity Filter | | | None |
| Quantification | Protein LFQ Method | | | Automatic |
| Quantification | Quantity MS Level | | | MS2 |
| Quantification | Quantity Type | | | Area |
| Quantification | Cross-Run Normalization | Normalization Filter Type | | None |
| Quantification | Cross-Run Normalization | Normalization Strategy | | Automatic |
| Quantification | Cross-Run Normalization | Row Selection | | Automatic |

*(continued)*

| Level 1 | Level 2 | Level 3 | Level 4 | Value |
|---|---|---|---|---|
| Quantification | Perform background noise removal | | | TRUE |
| Quantification | Quantification window | | | Not Synchronized (SN 17) |
| Quantification | Interference Correction | Only Identified Peptides | | TRUE |
| Quantification | Interference Correction | Exclude All Multi-Channel Interferences | | TRUE |
| Quantification | Interference Correction | MS1 Min | | 2 |
| Quantification | Interference Correction | MS2 Min | | 3 |
| Quantification | Major Group Quantity | | | Mean peptide quantity |
| Quantification | Minor (Peptide) Grouping | | | by Stripped Sequence |
| Quantification | Major (Protein) Grouping | | | by Protein Group Id |
| Quantification | Major Group Top N | Max | | 3 |
| Quantification | Major Group Top N | Min | | 1 |
| Quantification | Minor Group Quantity | | | Mean precursor quantity |
| Quantification | Minor Group Top N | Max | | 3 |
| Quantification | Minor Group Top N | Min | | 1 |
| Quantification | Use Log2 Quantity Filter | | | FALSE |
| Quantification | Perform IM Peak Picking for Quantification | | | TRUE |
| Workflow | Multi-Channel Workflow Definition | Fallback Option | | Labeled |
| Workflow | Profiling Strategy | | | None |
| Workflow | Hybrid (DDA + DIA) Library | | | FALSE |
| Workflow | In-Silico Library Optimization | | | FALSE |

*(continued)*

| Level 1 | Level 2 | Level 3 | Level 4 | Value |
|---|---|---|---|---|
| Workflow | Unify Peptide Peaks Strategy | | | None |
| XIC Extraction | XIC IM Extraction Window | Correction Factor | | 1 |
| XIC Extraction | XIC RT Extraction Window | Correction Factor | | 1 |
| XIC Extraction | MS1 Mass Tolerance Strategy | Tolerance (ppm) | | 7 |
| XIC Extraction | MS2 Mass Tolerance Strategy | Tolerance (ppm) | | 7 |

### 4 DIA-NN settings

The raw data analysis at the Utrecht site was conducted using DIA-NN (version 1.9). The analysis utilized and in-house blood protein database, that includes the SwissProt *Homo sapiens* sequences of the 2,445 proteins that are found in blood quantified, by either ELISA or MS, in blood according to the Human Protein Atlas (acquired November 13 2023). A detailed list of all parameters used is provided in the table below (Supplemental Table 5).

**Supplemental Table 5:** DIA-NN settings

| parameter | value |
|---|---|
| Cross-run normalization | RT-dependent |
| Deep learning-based spectra, RTs and Ims prediction | enabled |
| FASTA digest for library-free search | enabled |
| Fixed modifications | N-term M excision, C carbamidomethylation |
| Fragment ion m/z range | 200-1800 |
| Heuristic protein inference | disabled |
| Library generation | Ids, RT & IM profiling |
| Mass accuracy | 20 |
| Maximum number of variable modifications | 1 |
| MBR | enabled |
| Missed cleavages | 2 |
| MS1 accuracy | 10 |

*(continued)*

| parameter | value |
|---|---|
| Neural network classifier | Single-pass mode |
| No shared spectra | enabled |
| Peptide length range | 7-32 |
| Peptideoforms | enabled |
| Precursor charge range | 1-4 |
| Precursor m/z range | 300-1800 |
| Protein inference | Genes |
| Quantification strategy | Legacy (direct) |
| Threads | 40 |
| Unrelated runs | enabled |
| Variable modifications | Oxidation (M) |

### 5 used R packages

The data analysis was carried out using R and the listed packages below (Supplemental Table 6).

**Supplemental Table 6:** R packages used for the data analysis

| Name | Version | Citation |
|---|---|---|
| base | 4.4.1 | R: A Language and Environment for Statistical Computing 2024 |
| broom | 1.0.10 | broom: Convert Statistical Objects into Tidy Tibbles 2025 |
| dendextend | 1.19.1 | dendextend: an R package for visualizing, adjusting, and comparing trees of hierarchical clustering 2015 |
| factoextra | 1.0.7 | factoextra: Extract and Visualize the Results of Multivariate Data Analyses 2020 |
| FactoMineR | 2.12 | FactoMineR: A Package for Multivariate Analysis 2008 |
| furrr | 0.3.1 | furrr: Apply Mapping Functions in Parallel using Futures 2022 |
| fuzzyjoin | 0.1.6.1 | fuzzyjoin: Join Tables Together on Inexact Matching 2025 |
| ggh4x | 0.3.1 | ggh4x: Hacks for 'ggplot2' 2025 |
| gghalves | 0.1.4 | gghalves: Compose Half-Half Plots Using Your Favourite Geoms 2022 |
| ggrepel | 0.9.6 | ggrepel: Automatically Position Non-Overlapping Text Labels with 'ggplot2' 2024 |
| ggsignif | 0.6.4 | ggsignif: R Package for Displaying Significance Brackets for 'ggplot2' 2021 |
| ggstatsplot | 0.13.3 | Visualizations with statistical details: The 'ggstatsplot' approach 2021 |
| ggtext | 0.1.2 | ggtext: Improved Text Rendering Support for 'ggplot2' 2022 |
| glue | 1.8.0 | glue: Interpreted String Literals 2024 |
| helfRlein | 1.5.0 | helfRlein: R Helper Functions 2024 |
| kernlab | 0.9.33 | kernlab: Kernel-Based Machine Learning Lab 2024 |
| paletteer | 1.6.0 | paletteer: Comprehensive Collection of Color Palettes 2021 |
| patchwork | 1.3.2 | patchwork: The Composer of Plots 2025 |
| psych | 2.5.6 | psych: Procedures for Psychological, Psychometric, and Personality Research 2025 |
| purrr | 1.1.0 | purrr: Functional Programming Tools 2025 |

*(continued)*

| Name | Version | Citation |
|---|---|---|
| RColorBrewer | 1.1.3 | RColorBrewer: ColorBrewer Palettes 2022 |
| readxl | 1.4.5 | readxl: Read Excel Files 2025 |
| rsample | 1.3.1 | rsample: General Resampling Infrastructure 2025 |
| scales | 1.4.0 | scales: Scale Functions for Visualization 2025 |
| SpectroPipeR | 0.4.4 | SpectroPipeR — a streamlining post Spectronaut® DIA-MS data analysis R package 2025 |
| tidymodels | 1.4.1 | Tidymodels: a collection of packages for modeling and machine learning using tidyverse principles. 2020 |
| tidyverse | 2.0.0 | Welcome to the tidyverse 2019 |
| umap | 0.2.10.0 | umap: Uniform Manifold Approximation and Projection 2023 |
| yardstick | 1.3.2 | yardstick: Tidy Characterizations of Model Performance 2025 |

### 6 Plama Proteome Profiling

To evaluate the quality of the plasma samples the methodology from Geyer *et al* (2019) was used. Using the MaxLFQ protein intensities, proportions were calculated for platelets, erythrocytes, and coagulation contamination using their marker panels. This approach showed that only a marginal number of samples possess higher contaminations for platelets, erythrocytes, or coagulation (Supplemental Figure 1) and that the overall sample quality of the cohort is in an acceptable range.

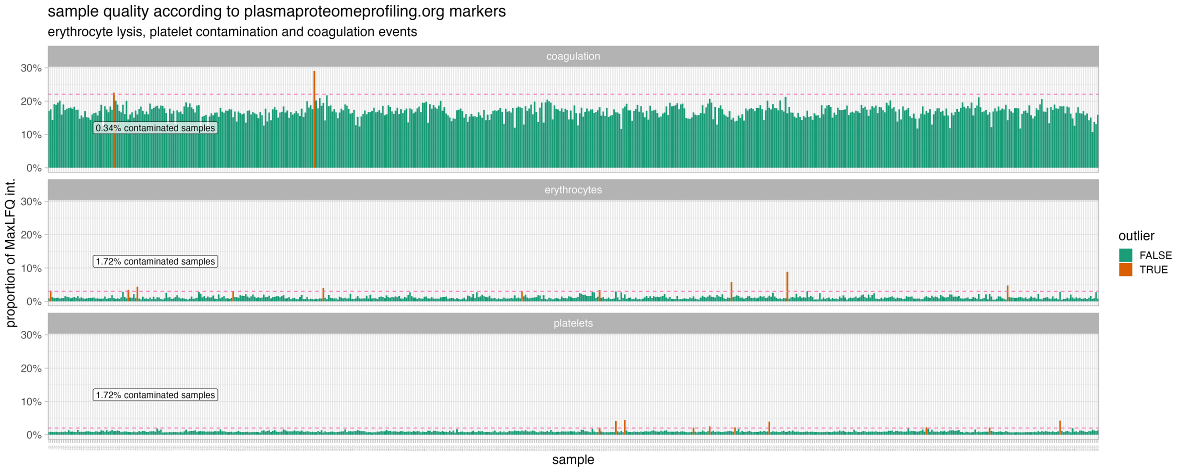

**Supplemental Figure 1:** Identification of problematic plasma samples due to sampling and effective quality control of the samples using the PlasmaProteomeProfiling gene set list. The PlasmaProteomeProfiling marker list was used to compute MaxLFQ protein proportions per sample, and outliers exceeding three standard deviations above the harmonic mean were flagged.

### 7 R-squared of linear modeling

The log10-scaled protein intensities were used to determine the absolute concentrations of proteins. This was done by fitting a linear model to the log10-scaled reference plasma protein concentrations measured by Gaither *et al* (2020). The model was tested for different combinations, including iBAQ ~ g/l, iBAQ ~ nM, MaxLFQ ~ g/l, and MaxLFQ ~ nM. For the Greifswald site, the iBAQ~nM model showed the best performance (Supplemental Figure 2), whereas for the Utrecht site the MaxLFQ~g/l and MaxLFQ~nM performed the equally better. We proceeded with the MaxLFQ~nM model, to achieve better comparisons with the Greifswald site.

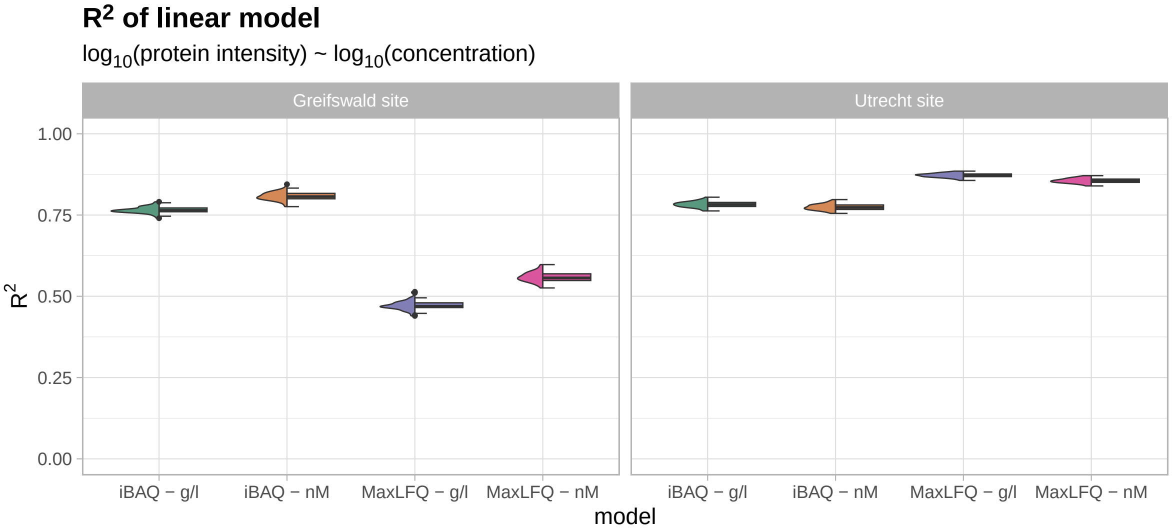

**Supplemental Figure 2:** R-squared of linear modeling of absolute protein concentrations (either g/L or nM) against relative protein intensities (iBAQ or MaxLFQ) across different acquisition sites.

### 8 multi-class sample prediction

To determine the feasibility of reattributing a random sample from a donor to its specific donor using the quantitative plasma protein profile, we employed a leave-one-out multi-class support vector machine (SVM) modeling approach (Supplemental Figure 3). Therefore, missing values were imputed by randomly sampling from the lower 5th percentile of observed concentrations. The predictors were standardized to maintain consistency across the analysis, and a fixed cost parameter (C = 1) was applied. After 1000 iterations probabilistic predictions were generated, and performance metrics were computed. The SVM achieved an accuracy of 0.98 in the median, indicating that when using a blinded random single sample with extremely high certainty, it can be accurately assigned to a donor.

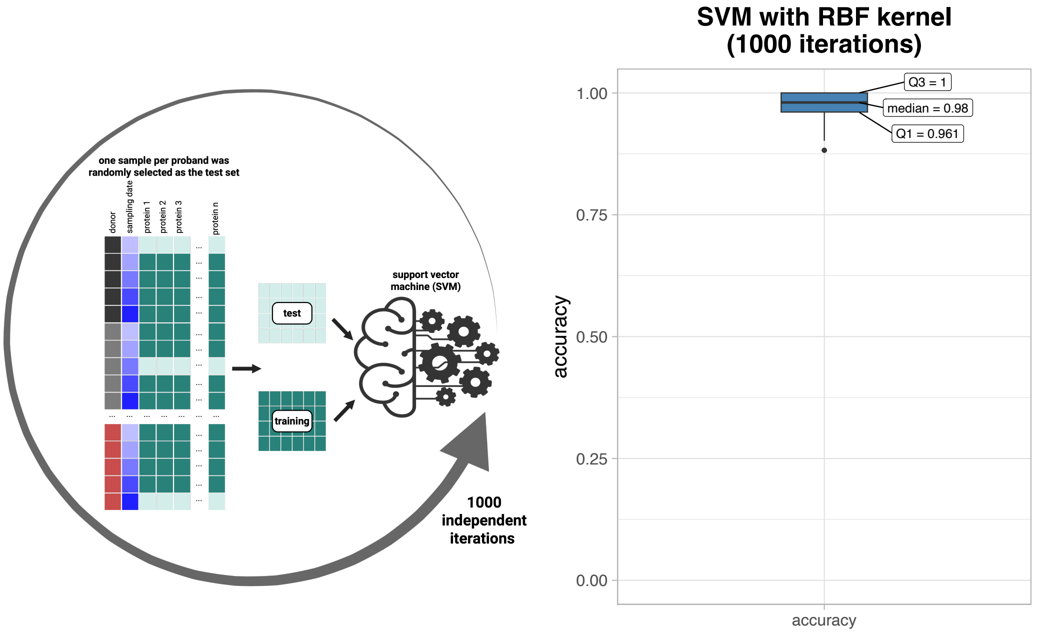

**Supplemental Figure 3:** Results of the multi-class SVM with RBF kernel modeling. Accuracy is depicted in a boxplot.

### 9 immunoglobulin sub-types and CD5L

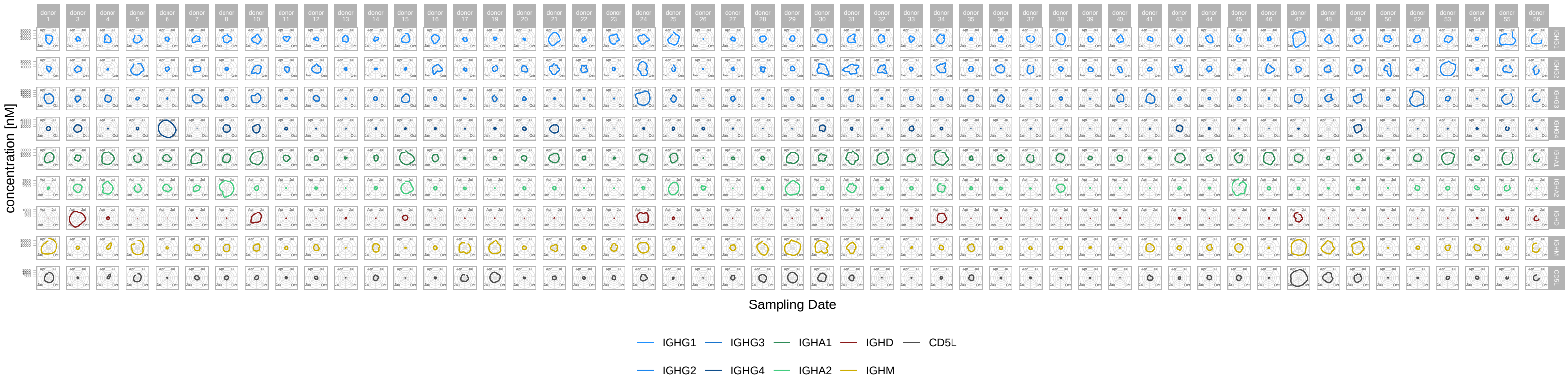

**Supplemental Figure 4:** Nanomolar (nM) concentrations of immunoglobulin subclasses (IgG, IgA, IgM, IgD) and CD5L are visualized per donor using radar plots. In this representation, a smaller circular diameter indicates a lower relative concentration compared to other donors. The closer the plotted line approximates a circle, the more stable the concentration remains over the 12-month observation period.

### 10 single plots IgG, IgA subtypes and IgM, IgD and CD5L

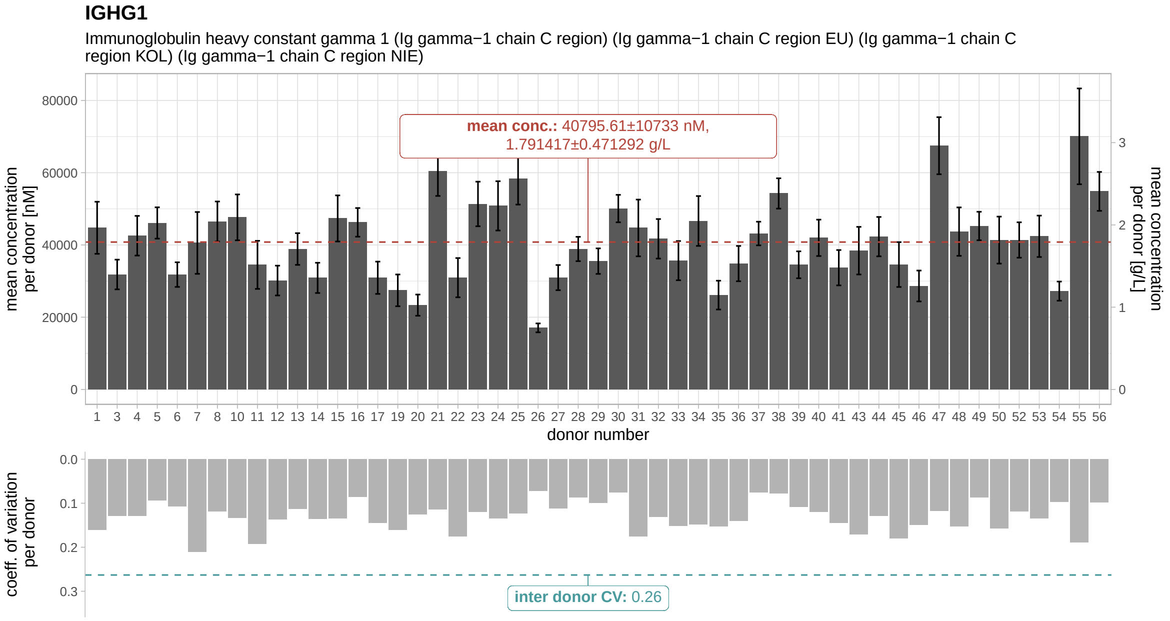

**Supplemental Figure 5:** Bar plot of IgG1 concentrations - The bar plot illustrates the mean and standard deviation of IgG1 concentrations across time points for each individual. Corresponding values in grams per liter (g/L) were derived from nanomolar (nM) concentrations using molecular weight-based conversion. The mean (dashed line) and standard deviation across donors are indicated in the label box. The barplot below presents the coefficient of variation (CV) across time points per donor. The inter-donor coefficient of variation is depicted as a dashed turquoise line, and its exact value is provided in the label.

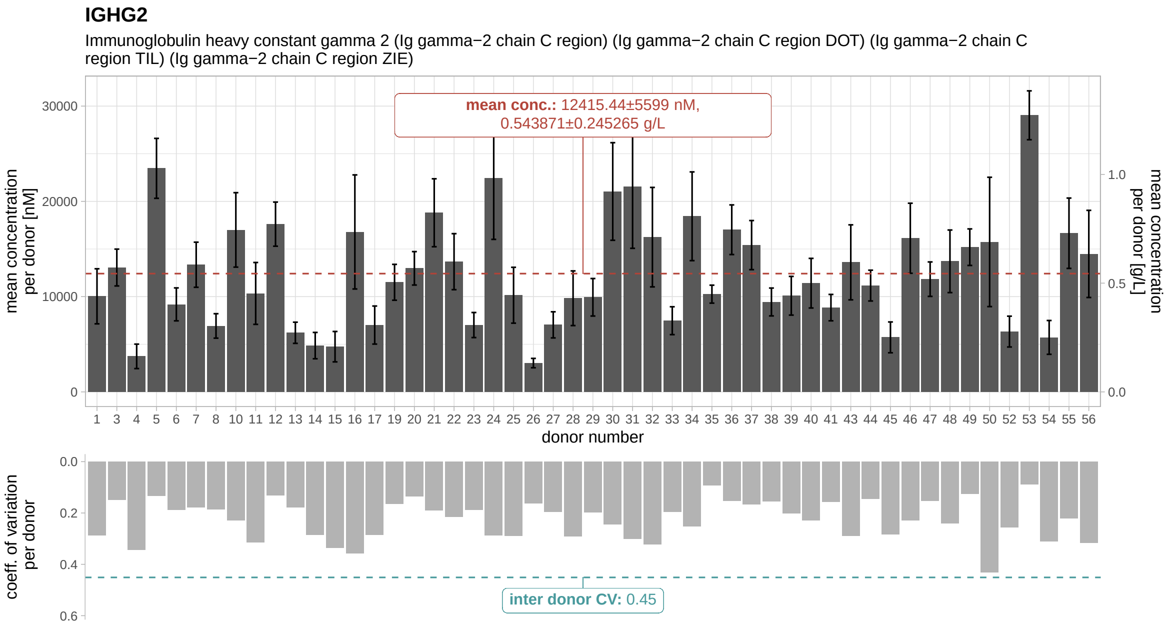

**Supplemental Figure 6:** Bar plot of IgG2 concentrations - The bar plot illustrates the mean and standard deviation of IgG2 concentrations across time points for each individual. Corresponding values in grams per liter (g/L) were derived from nanomolar (nM) concentrations using molecular weight-based conversion. The mean (dashed line) and standard deviation across donors are indicated in the label box. The barplot below presents the coefficient of variation (CV) across time points per donor. The inter-donor coefficient of variation is depicted as a dashed turquoise line, and its exact value is provided in the label.

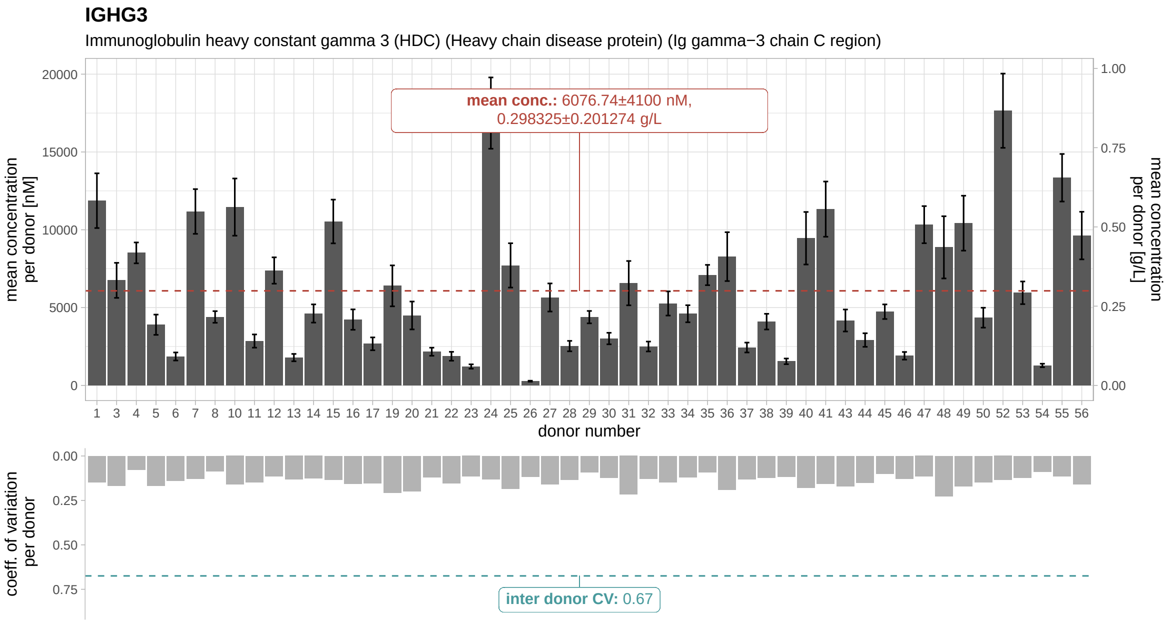

**Supplemental Figure 7:** Bar plot of IgG3 concentrations - The bar plot illustrates the mean and standard deviation of IgG3 concentrations across time points for each individual. Corresponding values in grams per liter (g/L) were derived from nanomolar (nM) concentrations using molecular weight-based conversion. The mean (dashed line) and standard deviation across donors are indicated in the label box. The barplot below presents the coefficient of variation (CV) across time points per donor. The inter-donor coefficient of variation is depicted as a dashed turquoise line, and its exact value is provided in the label.

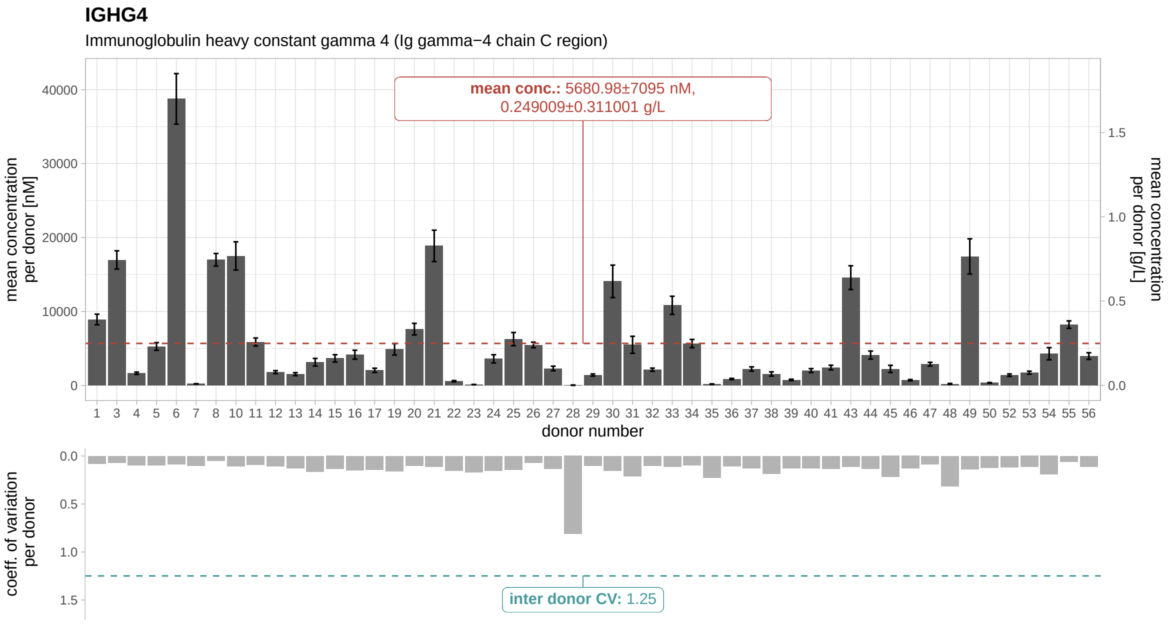

**Supplemental Figure 8:** Bar plot of IgG4 concentrations - The bar plot illustrates the mean and standard deviation of IgG4 concentrations across time points for each individual. Corresponding values in grams per liter (g/L) were derived from nanomolar (nM) concentrations using molecular weight-based conversion. The mean (dashed line) and standard deviation across donors are indicated in the label box. The barplot below presents the coefficient of variation (CV) across time points per donor. The inter-donor coefficient of variation is depicted as a dashed turquoise line, and its exact value is provided in the label.

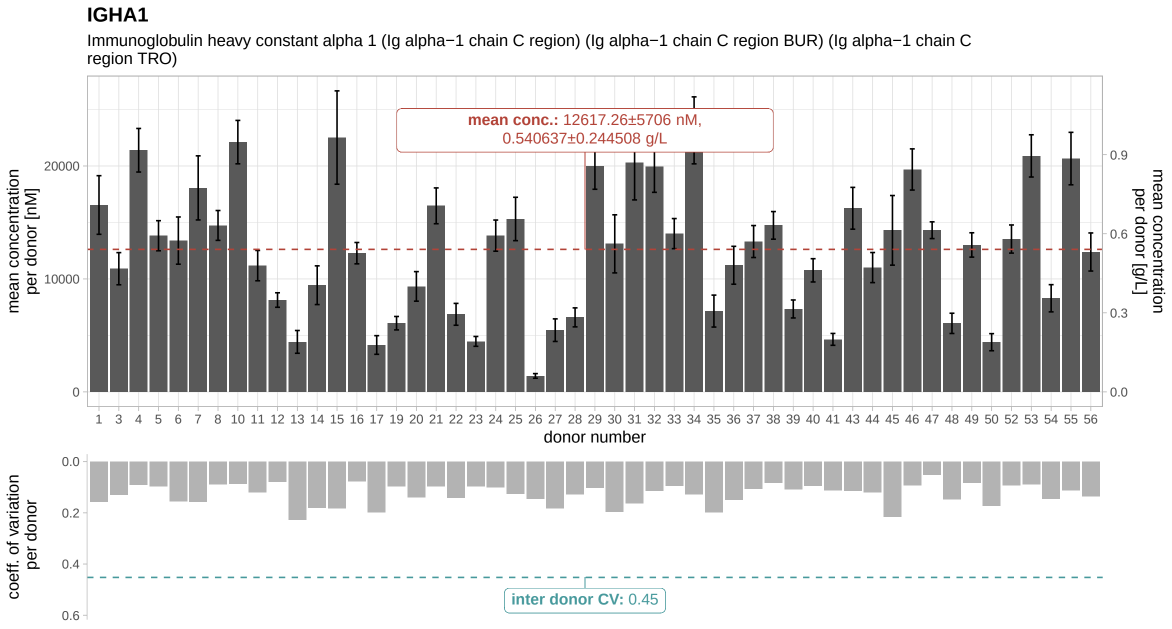

**Supplemental Figure 9:** Bar plot of IgA1 concentrations - The bar plot illustrates the mean and standard deviation of IgA1 concentrations across time points for each individual. Corresponding values in grams per liter (g/L) were derived from nanomolar (nM) concentrations using molecular weight-based conversion. The mean (dashed line) and standard deviation across donors are indicated in the label box. The barplot below presents the coefficient of variation (CV) across time points per donor. The inter-donor coefficient of variation is depicted as a dashed turquoise line, and its exact value is provided in the label.

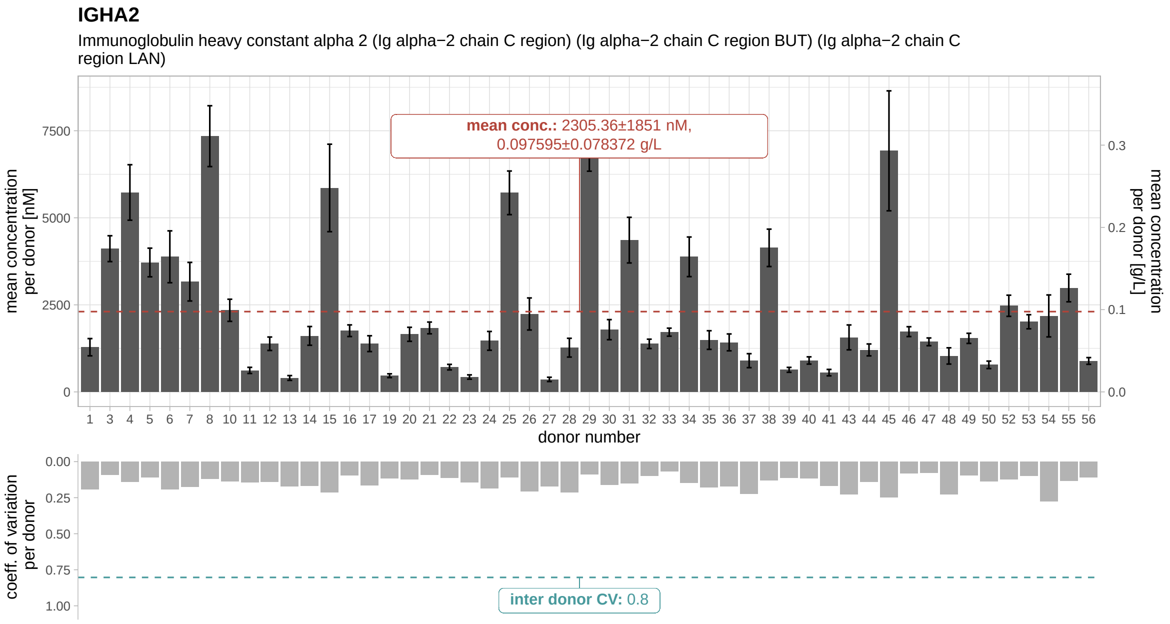

**Supplemental Figure 10:** Bar plot of IgA2 concentrations - The bar plot illustrates the mean and standard deviation of IgA2 concentrations across time points for each individual. Corresponding values in grams per liter (g/L) were derived from nanomolar (nM) concentrations using molecular weight-based conversion. The mean (dashed line) and standard deviation across donors are indicated in the label box. The barplot below presents the coefficient of variation (CV) across time points per donor. The inter-donor coefficient of variation is depicted as a dashed turquoise line, and its exact value is provided in the label.

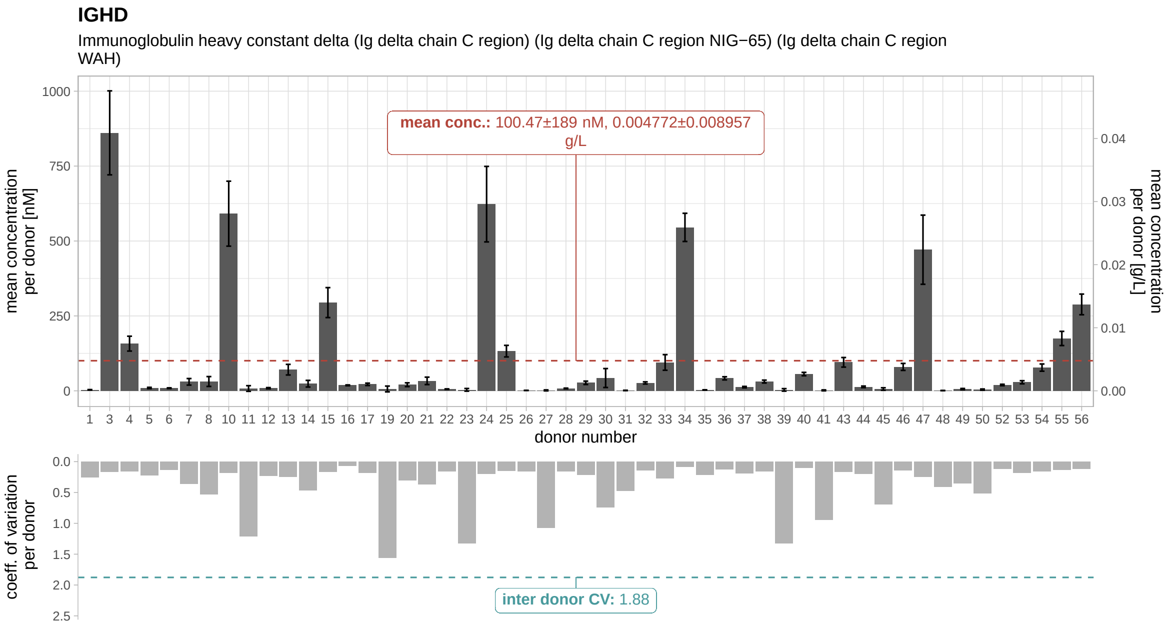

**Supplemental Figure 11:** Bar plot of IgD concentrations - The bar plot illustrates the mean and standard deviation of IgD concentrations across time points for each individual. Corresponding values in grams per liter (g/L) were derived from nanomolar (nM) concentrations using molecular weight-based conversion. The mean (dashed line) and standard deviation across donors are indicated in the label box. The barplot below presents the coefficient of variation (CV) across time points per donor. The inter-donor coefficient of variation is depicted as a dashed turquoise line, and its exact value is provided in the label.

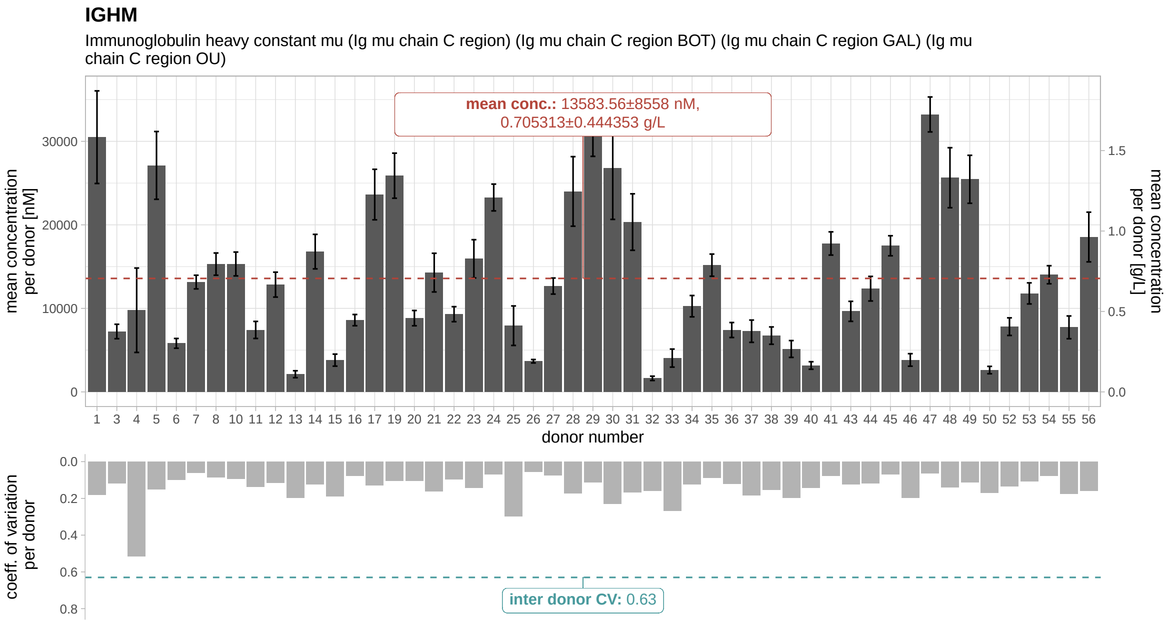

**Supplemental Figure 12:** Bar plot of IgM concentrations - The bar plot illustrates the mean and standard deviation of IgM concentrations across time points for each individual. Corresponding values in grams per liter (g/L) were derived from nanomolar (nM) concentrations using molecular weight-based conversion. The mean (dashed line) and standard deviation across donors are indicated in the label box. The barplot below presents the coefficient of variation (CV) across time points per donor. The inter-donor coefficient of variation is depicted as a dashed turquoise line, and its exact value is provided in the label.

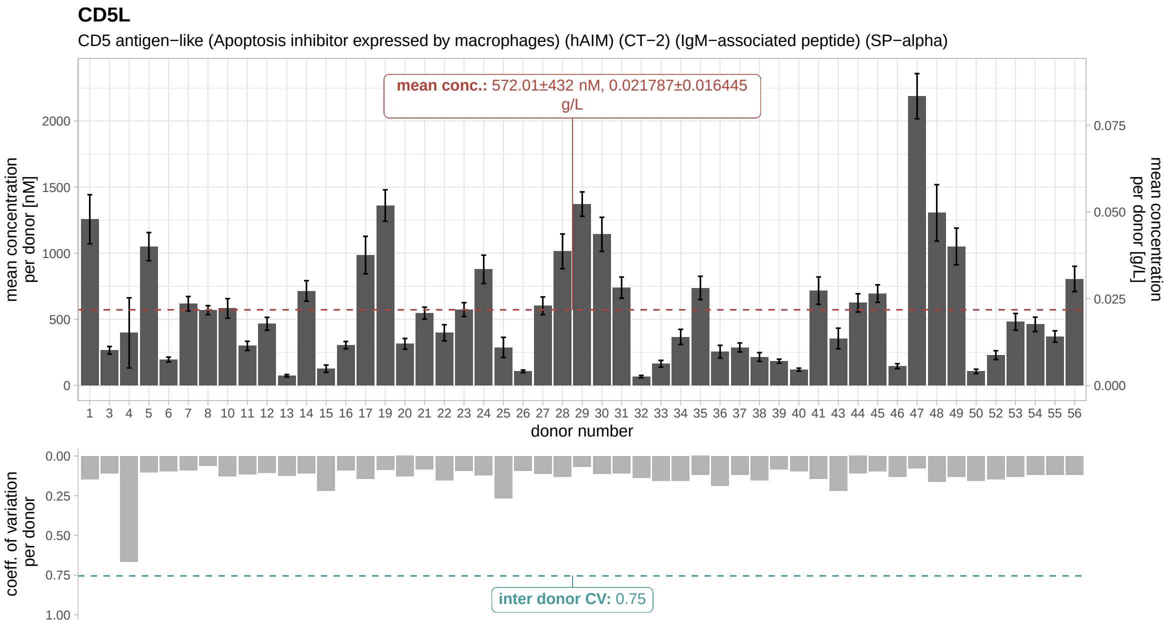

**Supplemental Figure 13:** Bar plot of CD5L concentrations - The bar plot illustrates the mean and standard deviation of CD5L concentrations across time points for each individual. Corresponding values in grams per liter (g/L) were derived from nanomolar (nM) concentrations using molecular weight-based conversion. The mean (dashed line) and standard deviation across donors are indicated in the label box. The barplot below presents the coefficient of variation (CV) across time points per donor. The inter-donor coefficient of variation is depicted as a dashed turquoise line, and its exact value is provided in the label.

### 11 IgG subclass distribution

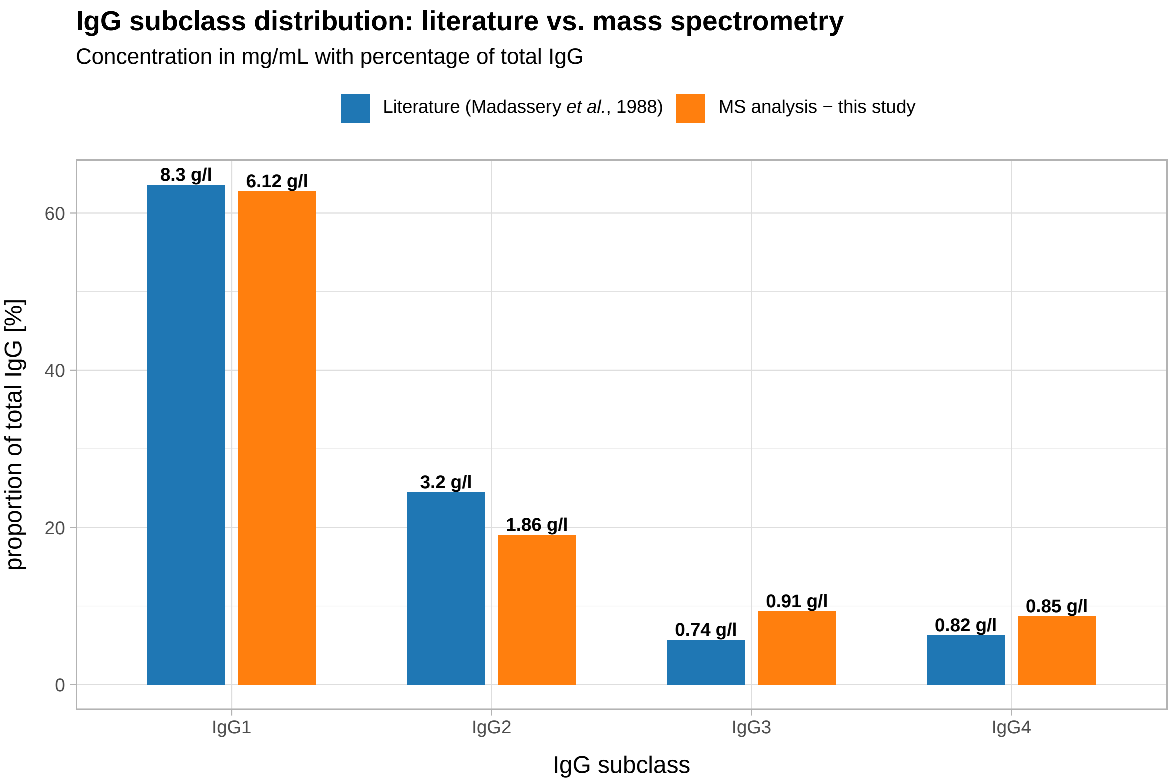

**Supplemental Figure 14:** Bar plot depicts the proportion of IgG subclasses known from literature and measured in in the TIMES cohort via mass spectroemtry

### 12 selected inflammation markers

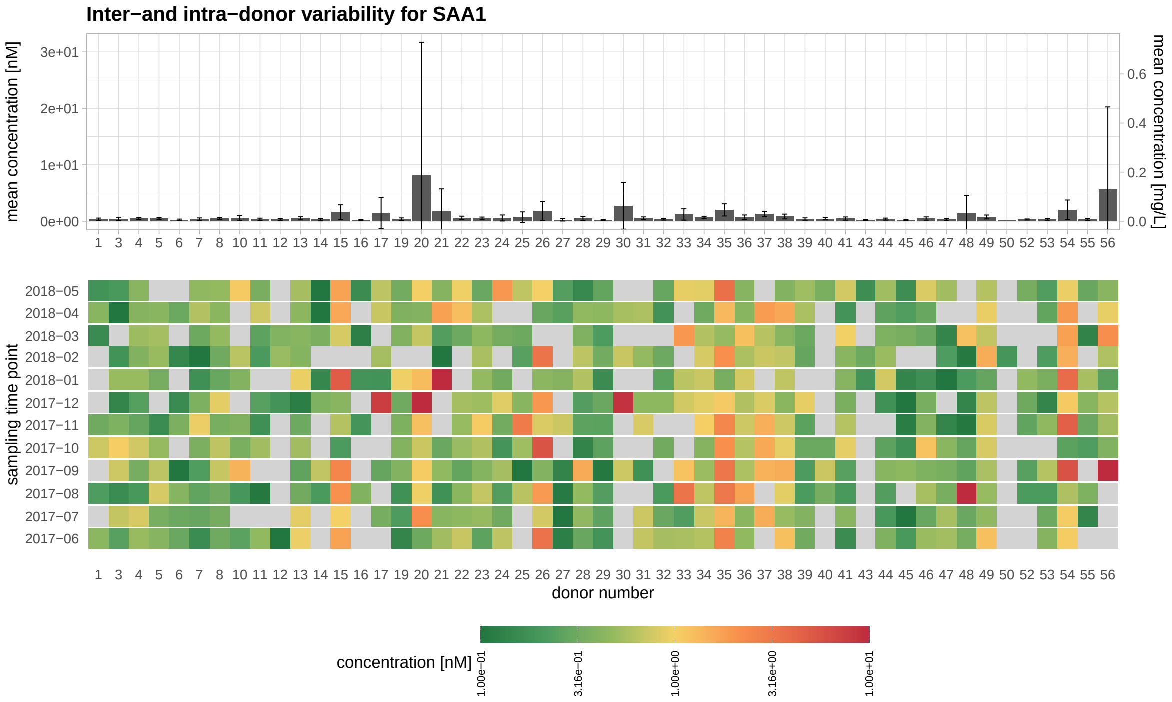

**Supplemental Figure 15:** SAA1 mean concentration shown in barplot (top panel). The tile plot below (lower panel) depicts the temporal profiles of SAA1 in donors (grey color represent missing values).

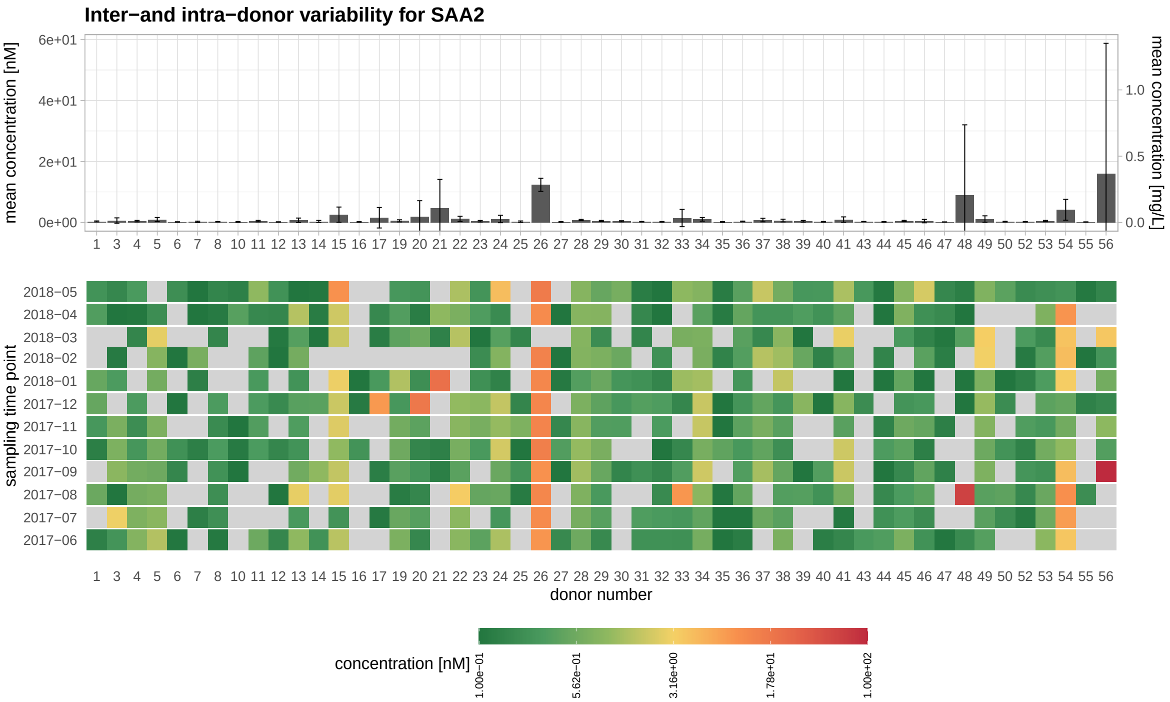

**Supplemental Figure 16:** SAA2 mean concentration shown in barplot (top panel). The tile plot below (lower panel) depicts the temporal profiles of SAA2 in donors (grey color represent missing values).

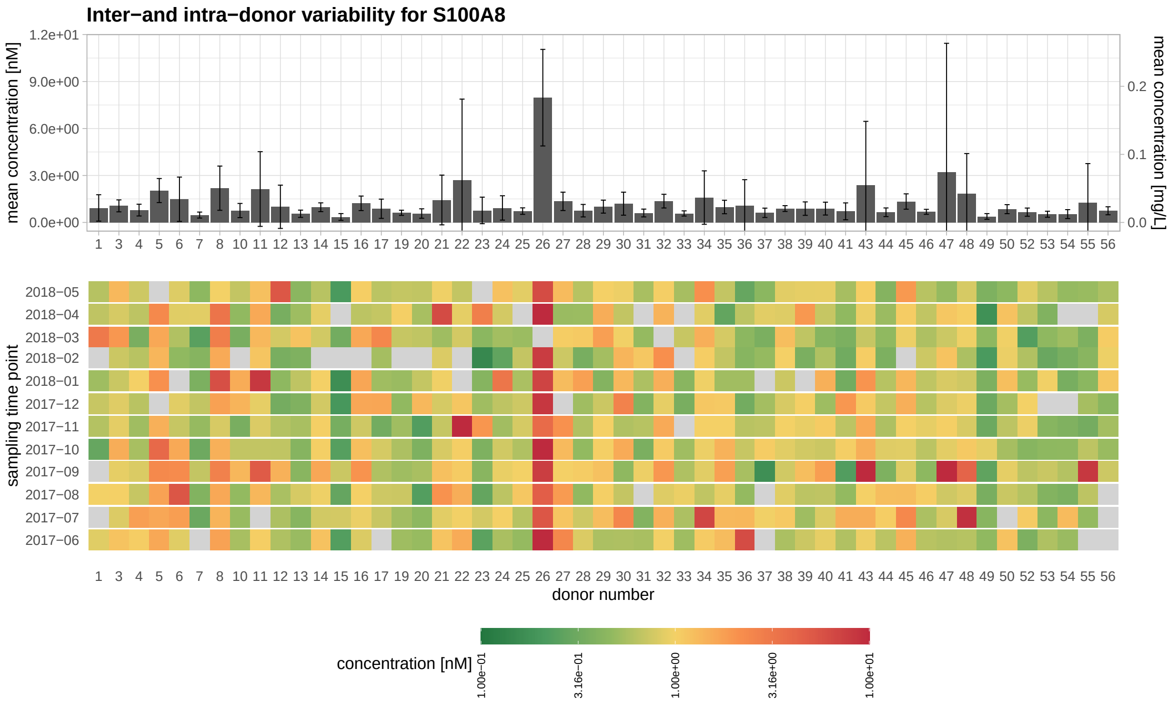

**Supplemental Figure 17:** S100A8 mean concentration shown in barplot (top panel). The tile plot below (lower panel) depicts the temporal profiles of S100A8 in donors (grey color represent missing values).

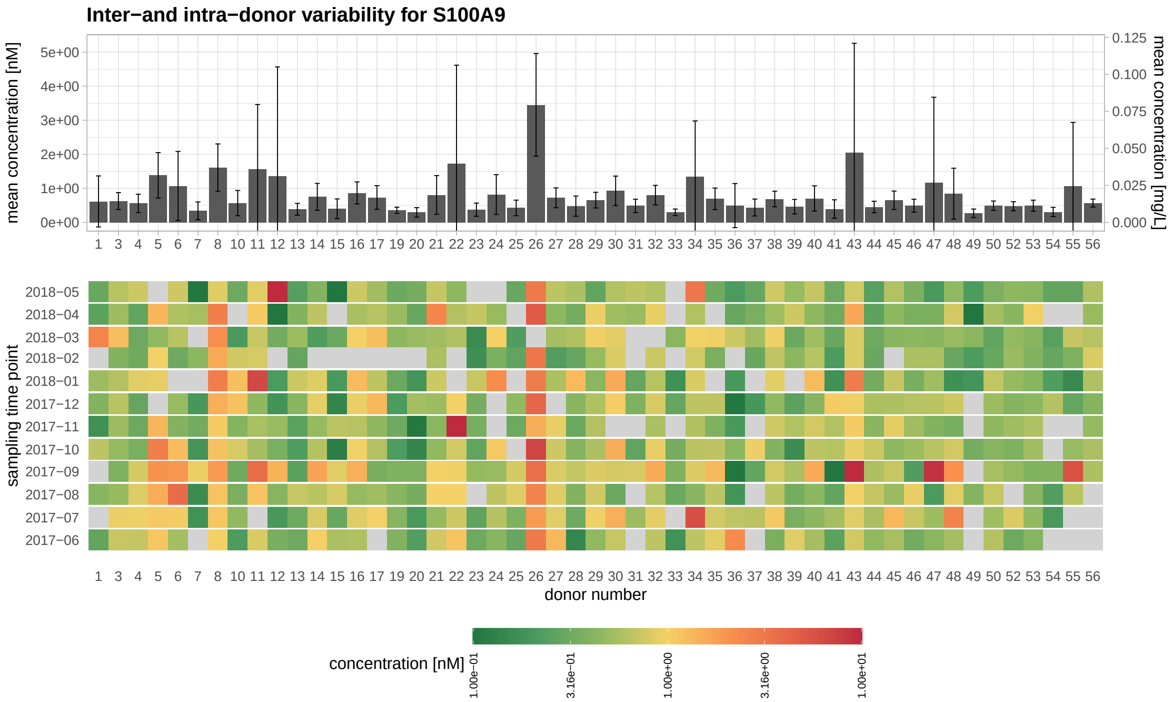

**Supplemental Figure 18:** S100A9 mean concentration shown in barplot (top panel). The tile plot below (lower panel) depicts the temporal profiles of S100A9 in donors (grey color represent missing values).

### References

Gaither C, Popp R, Mohammed Y & Borchers CH (2020) Determination of the concentration range for 267 proteins from 21 lots of commercial human plasma using highly multiplexed multiple reaction monitoring mass spectrometry. *Analyst* 145: 3634–3644

Geyer PE, Voytik E, Treit PV, Doll S, Kleinhempel A, Niu L, Müller JB, Buchholtz M, Bader JM, Teupser D, *et al* (2019) Plasma proteome profiling to detect and avoid sample-related biases in biomarker studies. *EMBO Molecular Medicine* 11: e10427
